# The Mediator CDK8-Cyclin C complex modulates vein patterning in *Drosophila* by stimulating Mad-dependent transcription

**DOI:** 10.1101/360628

**Authors:** Xiao Li, Mengmeng Liu, Xingjie Ren, Nicolas Loncle, Qun Wang, Rajitha-Udakara-Sampath Hemba-Waduge, Muriel Boube, Henri-Marc G. Bourbon, Jian-Quan Ni, Jun-Yuan Ji

**Author notes:** Current address: Femtera Laboratories LLC., Indianapolis, IN 46202, United States of America. Corresponding author: Department of Molecular and Cellular Medicine, Texas A&M University Health Science Center, College Station, TX 77843.

## Abstract

Dysregulations of CDK8 and its regulatory partner CycC, two subunits of the conserved Mediator complex, have been linked to diverse human diseases such as cancer, thus it is essential to understand the regulatory network mobilizing the CDK8-CycC complex in both normal development and tumorigenesis. To identify upstream regulators or downstream effectors of CDK8, we performed a dominant modifier genetic screen in *Drosophila* based on the defects in vein patterning caused by specific depletion or overexpression of CDK8 or CycC in wing imaginal discs. We identified 26 genomic loci whose haploinsufficiency can modify these CDK8-specific phenotypes. Further analysis of two deficiency lines and mutant alleles led us to identify interactions between CDK8-CycC and the components of the Decapentaplegic (Dpp, the *Drosophila* homolog of TGFβ) signaling pathway. We observed that CDK8-CycC positively regulates transcription activated by Mad (Mothers against dpp), the primary transcription factor downstream of the Dpp/TGFβ signaling pathway. CDK8 can directly interact with Mad *in vitro* through the linker region between the DNA-binding MH1 (Mad homology 1) domain and the carboxy terminal MH2 transactivation domain. Besides CDK8 and CycC, further analyses of other subunits of the Mediator complex have revealed six additional Mediator subunits that are required for Mad-dependent transcription in the wing discs, including Med12, Med13, Med15, Med23, Med24, and Med31. Furthermore, CDK9 and Yorkie also positively regulate Mad-dependent gene expression *in vivo*. These results suggest that the Mediator complex may coordinate with other transcription cofactors in regulating Mad-dependent transcription during the wing vein patterning in *Drosophila*.

**Significance:** CDK8 is a conserved subunit of the transcription cofactor Mediator complex that bridges transcription factors with RNA Polymerase II in eukaryotes. Here we explore the role of CDK8 in *Drosophila* by performing a dominant modifier genetic screen based on vein patterning defects caused by alteration of CDK8-specific activities. We show that components of the Dpp/TGFβ signaling pathway genetically interact with CDK8; CDK8 positively regulates gene expression activated by Mad, the key transcription factor downstream of Dpp/TGFβ signaling, by directly interacting with the linker region of Mad protein. Given the fundamental roles of Dpp/TGFβ signaling in regulating development and its misregulation in various diseases, understanding how Mad/Smad interacts the Mediator complex may have broad implications in understanding and treating these diseases.

## Introduction

Composed of up to 30 conserved subunits, the transcription cofactor Mediator complex plays critical roles in modulating RNA polymerase II (Pol II)-dependent gene expression by functioning as a molecular bridge linking transcriptional activators and the general transcription machinery in almost all eukaryotes (1-5). Biochemical purification of the human Mediator complexes has revealed the CDK8 (Cyclin-Dependent Kinase 8) module, composed of CDK8 (or its paralogue CDK19, also known as CDK8L), Cyclin C (CycC), Med12 (or Med12L), and Med13 (or Med13L), and the small Mediator complex, composed of 26 subunits that are divided into the head, middle, and tail modules (6-9). CDK8 is the only Mediator subunit with enzymatic activities. The CDK8 module (also known as CKM, for CDK8 kinase module) has been proposed to function in two modes: it can reversibly bind with the small Mediator complex to form the large Mediator complex, thereby physically blocking the interaction between the small Mediator complex and the general transcription machinery, notably with RNA Pol II itself; and alternatively, CDK8 can function as a kinase that phosphorylates different substrates, particularly transcriptional activators such as E2F1 (10, 11), Notch-ICD (intracellular domain of Notch) (12), p53 (13), Smad proteins (14, 15), SREBP (sterol regulatory element-binding protein) (16), and STAT1 (signal transducer and activator of transcription 1) (17). These characterized functions of CDK8 highlight fundamental roles of CKM in regulating transcription.

Besides its roles in specific developmental and physiological contexts, the CKM subunits are dysregulated in a variety of human diseases, such as cancers (18-21). For example, CDK8 has been reported to act as an oncoprotein in melanoma and colorectal cancers (22, 23). CDK8 and CDK19 are overexpressed in invasive ductal carcinomas, correlating with shorter relapse-free survival in breast cancer (24). Gain or amplification of CDK8 activity is sufficient to drive tumorigenesis in colorectal and pancreatic cancers in human, as well as skin cancer in fish (14, 22, 25-27). Because of these discoveries, there is a heightened interest in developing drugs targeting the CDK8 kinase for cancer treatment in recent years (28, 29). However, exactly how CDK8 dysregulation contributes to tumorigenesis remains poorly understood. The key is to reveal the function and regulation of CDK8 activity in different developmental and physiological processes.

The major bottleneck for addressing these critical gaps in our knowledge is the lack of *in vivo* readouts for CDK8-specific activities in metazoans. We overcame this challenge by generating tissue-specific phenotypes caused by varying CDK8 activities in *Drosophila*. After validating the specificity of these phenotypes using genetic, molecular, and cell biological approaches, we have performed a dominant modifier genetic screen to identify factors that interact with CDK8 *in vivo* based on these unique readouts for CDK8-specific activities. From the screen, we identified *Dad* (*Daughters against dpp*), encoding an inhibitory Smad in the Dpp (Decapentaplegic)/TGFβ (Transforming Growth Factor β) signaling pathway, as well as additional components of the Dpp signaling pathway, including *dpp, tkv* (*thickveins*, encoding the type I receptor for Dpp), *Mad* (*Mothers against dpp*) and *Medea*, encoding the Smad1/5 and Smad4 homologs, respectively, in *Drosophila*. Consistent to the previous biochemical analyses suggesting that CDK8 can phosphorylate *Drosophila* Mad or conserved sites in human Smad1 thereby stimulating their transcriptional activities (14, 15, 30), our results suggest that this regulatory mechanism *in vivo* is conserved in evolution. Furthermore, our analyses have revealed additional Mediator subunits and kinases involved in regulating the Mad/Smad-dependent transcription. These results, together with the previous studies, suggest that concerted recruitment of the Mediator complexes and other cofactors plays a pivotal role in regulating the Mad/Smad-dependent gene expression, a critical process for TGFβ signaling to function in a variety of biological and pathological contexts.

## Results

### Wing vein patterning phenotypes of altering CDK8 or CycC

To study the function and regulation of CDK8 and CycC *in vivo*, we have generated transgenic lines to either deplete them by RNA interference (RNAi) or conditionally overexpress the wild-type CDK8 using the Gal4-UAS system (see Materials and Methods for details). Normal *Drosophila* wings display stereotypical vein patterns, consisting of six longitudinal veins, L1 to L6, and two crossveins, the anterior crossvein and the posterior crossvein (Fig. 1A). Knocking down of CDK8 using the *nubbin-Gal4* line (*nub-Gal4*), which is specifically expressed in the wing pouch area of the wing imaginal discs (31), results in the formation of ectopic veins in the intervein region, especially around L2 and L5 (Fig. 1B). Similar phenotypes were observed with the depletion of CycC (Fig. 1C), or both CDK8 and CycC (Fig. 1D). In contrast, overexpression of wild-type CDK8 (*UAS-Cdk8^+^*) disrupts the L3 and L4 veins, as well as the crossveins (Fig. 1E), opposite to the phenotypes caused by depleting CDK8, or CycC, or both. However, overexpression of a kinase-dead (KD) CDK8 form (*UAS-Cdk8^KD^*) using the same approach does not affect the vein patterns (Fig. 1F), suggesting that the effects of CDK8 on vein phenotypes are dependent on the kinase activity of CDK8. These observations show that CDK8 and CycC are involved in regulating the vein patterning in *Drosophila*.

**Fig. 1.**
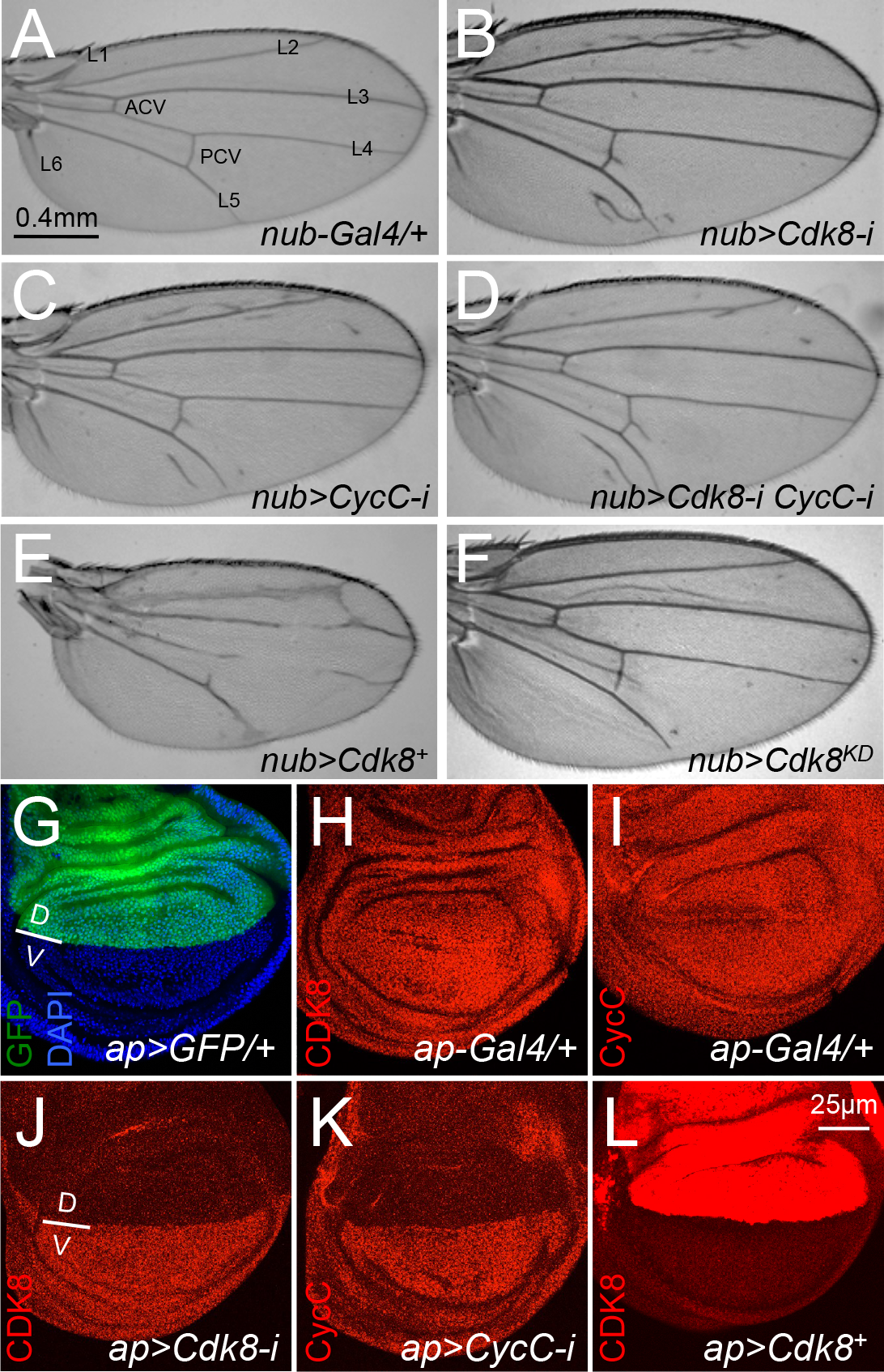
Wing vein patterning phenotypes caused by depletion or overexpression of CDK8 or CycC. Adult female wings of (A) *nub-Gal4/+* (control), note the longitudinal veins L1-L6, anterior crossvein (ACV), and posterior crossvein (PCV); (B) *w^1118^/+; nub-Gal4/+; UAS-Cdk8-RNAi/+*; (C) *w^1118^/+; nub-Gal4/+; UAS-CycC-RNAi/+*; (D) *w^1118^/+; nub-Gal4/+; UAS-Cdk8-RNAi CycC-RNAi/+*; (E) *w^1118^/+; nub-Gal4>UAS-Cdk8^+^/+*; and (F) *w^1118^/+; nub-Gal4/UASCdk8^KD^* under a 5X objective microscope. Confocal images of the wing pouch area of a L3 wondering larvae wing disc of (G) *ap-Gal4/UAS-2XGFP* with DAPI (blue) and GFP (green); (H) *ap-Gal4/+* with anti-CDK8 (red) staining; (I) *ap-Gal4/+* with anti-CycC (red) staining; (J) *ap-Gal4/+; UAS-Cdk8-RNAi/+* with anti-CDK8 (red) staining; (K) *ap-Gal4/+; UAS-CycC-RNAi/+* with anti-CycC (red) staining; and (L) *ap-Gal4/UAS-Cdk8^+^* with anti-CDK8 (red) staining. Note that the gain for confocal imaging in overexpression figures is lower than others to avoid over saturation of the signals. The dorsal/ventral boundary is shown in G. Scale bar in A (for A-F: 0.4mm), in L (for G-L): 25µm.

### Validation of the depletion or overexpression phenotype specificity

To verify the specificity of these phenotypes, we recombined the *nub-Gal4* line with the CDK8-RNAi, CycC-RNAi, or CDK8-overexpression lines, and then tested whether these vein phenotypes could be dominantly modified by *cdk8^K185^*, a null allele of *cdk8* (32). As shown in Fig. S1A, reducing CDK8 by half in ‘*cdk8^K185^/+*’ heterozygous background suppresses the vein defects caused by CDK8 overexpression. However, heterozygosity of *cdk8^K185^* does not further enhance the vein phenotype caused by CDK8-RNAi (Fig. S1B), indicating that RNAi of CDK8 may have depleted most of the CDK8 protein pool.

To further validate the specificity of the CDK8-directed phenotypes at the cellular level, we analyzed the protein levels of CDK8 and CycC in wing discs at the third instar wandering stage by immunostaining with CDK8 or CycC specific antibodies. For this, we used the *apterous-Gal4 (ap-Gal4)* line, which is specifically expressed within the dorsal compartment of the wing discs (Fig. 1G) (33), allowing us to use the ventral compartment of the same discs as the internal control. Normally, both the CDK8 (Fig. 1H) and CycC (Fig. 1I) proteins are uniformly distributed in the nuclei of all wing disc cells. Tissue-specific depletion of CDK8 (Fig. 2J), or CycC (Fig. 2K), or both (Figs. S1C, S1D) using the *ap-Gal4* line led to significantly reduced protein levels for CDK8 or CycC in the dorsal compartment. In contrast, overexpression of either wild-type (Fig. 1L) or kinase-dead (Fig. S1E) CDK8 driven by *ap-Gal4* specifically increased the levels of CDK8 protein in the dorsal compartment. Taken together, these genetic and cell biological analyses validate the specificity of both the antibodies and transgenic lines, demonstrating that these vein phenotypes are caused by specific gain or reduction of CDK8 activity *in vivo*.

### Identification of deficiency lines that can dominantly modify the vein phenotypes caused by varying CDK8

Based on these CDK8-specific vein phenotypes, we performed a dominant modifier genetic screen to identify gene products that can functionally interact with CDK8 *in vivo*. This approach has been successfully employed to reveal the regulatory networks for proteins of interest in *Drosophila* (34). The approach posits that if a protein interacts with CDK8-CycC *in vivo* in defining the vein patterns, then reducing its level by half may either enhance or suppress the sensitized vein phenotypes caused by specific alteration of the CDK8 activities. Accordingly, we can survey through the fly genome to search for factors that interact with CDK8-CycC by single genetic crosses.

To facilitate this screen approach, we generated three stocks with the following genotypes: “*w^1118^; nub-Gal4; UAS-Cdk8-RNAi*” (designated as “*nub>Cdk8-i*” for simplicity), “*w^1118^; nub-Gal4; UAS-CycC-RNAi*” (“*nub>CycC-i*”), and “*w^1118^; nub-Gal4, UAS-Cdk8^+^/CyO*” (“*nub>Cdk8^+^*”). We then conducted the primary screen by crossing these three lines in parallel with a collection of 490 deficiency (*Df*) lines (Table S1), which uncovers approximately 98% of the euchromatic genome. The vein patterns of the F1 females were inspected for enhancers and suppressors based on the following criteria: suppressors of the CDK8- or CycC-RNAi phenotypes would display fewer or no ectopic veins (e.g., Figs. 2A, 2C), while enhancers of the CDK8- or CycC-RNAi phenotypes would show extra ectopic veins (e.g., Fig. 2B, 2D). Similarly, the suppressors of the CDK8-overexpression phenotype are expected to have vein patterns similar to those of wild-type wings, particularly the L3/L4 (e.g., Fig. 2E), while enhancers of the CDK8-overexpression phenotype should further disrupt the vein patterns (e.g., Fig. 2F).

**Fig. 2.**
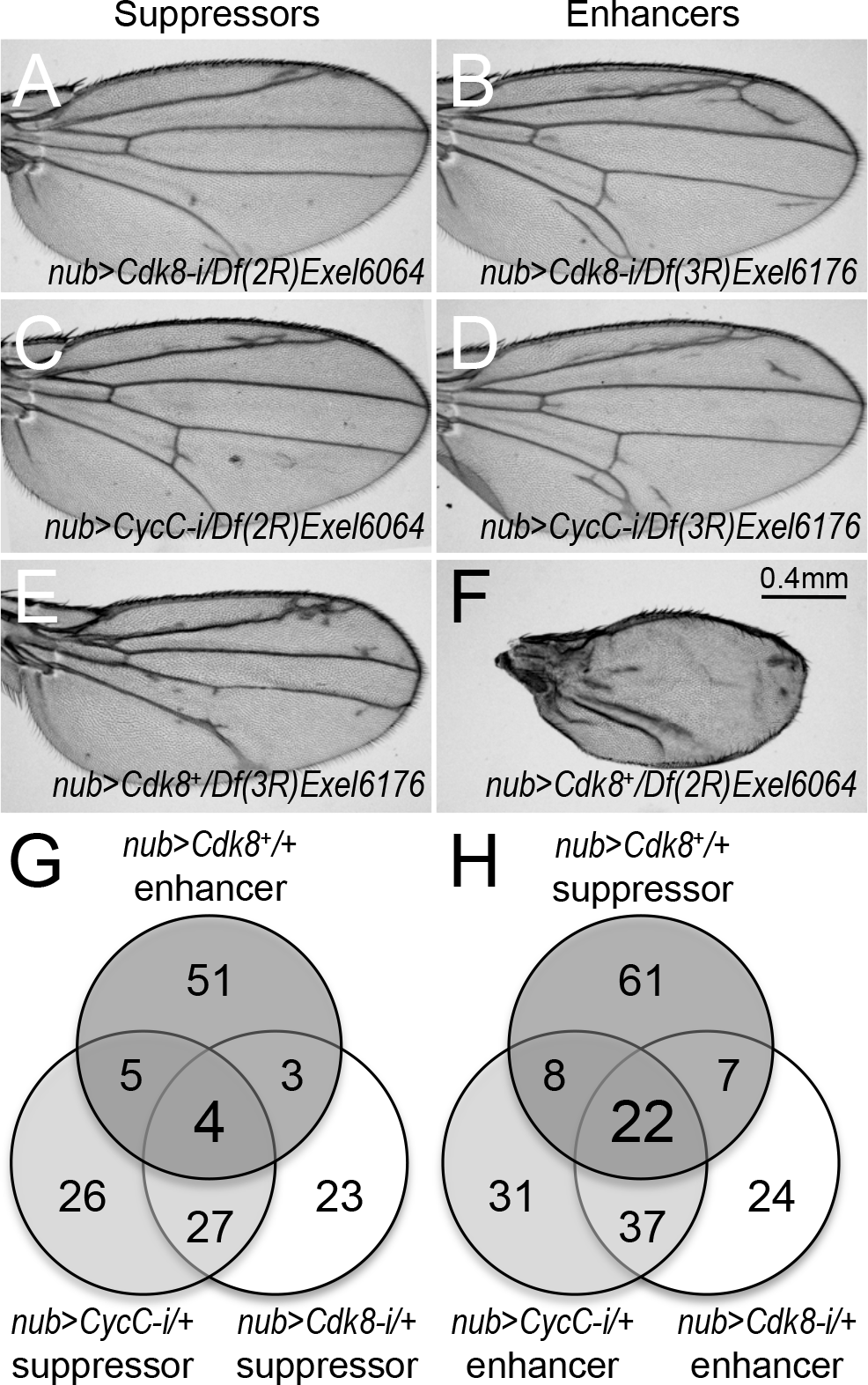
Identification of deficiency lines that can dominantly modify the vein phenotypes caused by altering CDK8 levels. Adult female wings of (A) *nub-Gal4/Df(2R)Exel6064; UASCdk8-RNAi*, an example of the CDK8 depletion phenotype suppressor lines; (B) *nub-Gal4/+; UAS-Cdk8-RNAi/ Df(3R)Exel6176*, an example of the CDK8 depletion phenotype enhancer lines; (C) *nub-Gal4/Df(2R)Exel6064; UAS-CycC-RNAi/+*, an example of the CycC depletion phenotype suppressor lines; (D) *nub-Gal4/+; UAS-CycC-RNAi/ Df(3R)Exel6176*, an example of the CycC depletion phenotype enhancer lines; (E) *nub-Gal4>UAS-Cdk8^+^/+; Df(3R)Exel6176 /+*, an example of the overexpression phenotypes suppressor lines; and (F) *nub-Gal4>UASCdk8^+^/Df(2R)Exel6064*, an example of the overexpression phenotypes enhancer lines. Scale bar in F: 0.4mm. (G and H) The number of suppressors and enhancers of the CDK8-CycC phenotypes as summarized using the Venn diagrams.

From these screens, we identified 57 suppressor and 90 enhancer *Df* lines for the CDK8-RNAi phenotype, and 62 suppressor and 98 enhancer *Df* lines for the CycC-RNAi phenotype. In addition, we identified 63 enhancer and 98 suppressor *Df* lines for the CDK8-overexpression phenotype (Fig. 2G, 2H). The results for all of these *Df* lines are summarized in Table S1. Of these dominant modifier *Df* lines, four of them suppressed the CDK8-RNAi and CycC-RNAi phenotypes but enhance the CDK8-overexpression phenotype (Fig. 2G, Table S2), while 22 of them enhance the CDK8-RNAi and CycC-RNAi phenotypes but suppress the CDK8-overexpression phenotype (Fig. 2H, Table S2). To further validate this genetic approach, we generated a transgenic line that allowed us to deplete both CDK8 and CycC simultaneously (“*w^1118^; nub-Gal4; UAS-Cdk8-RNAi, CycC-RNAi*”, referred to as “*nub>Cdk8-i CycC-i*”) with *nub-Gal4*, and observed the identical phenotypes to the ones caused by depleting either *Cdk8* or *CycC* alone (Fig. 1D). With the exception of one *Df* line, the rest of these 25 *Df* lines have consistently modified the ectopic vein phenotype caused by depletion of both CDK8 and CycC: four of the *Df* lines behaved as suppressors and 21 of them as enhancers (Table S2). These results show that the CDK8-specific vein phenotypes are modifiable and can be used to identify factors that functionally interact with CDK8 *in vivo*.

### Identification of *Dad* as an enhancer of the *nub>Cdk8i* and *nub>CycCi* phenotypes but a suppressor of the *Cdk8*-overexpression phenotype

To identify the specific genes uncovered by these dominant modifier *Df* lines, we analyzed these 25 genome regions with partial overlapping *Df* lines (Table S2). Interestingly, two partially overlapping *Df* lines, *Df(3R)BSC748* and *Df(3R)Exel6176*, enhanced the CDK8-RNAi and CycC-RNAi phenotypes, but suppressed the CDK8-overexpression phenotype (Fig. 3A). The overlapping region uncovers one specific gene, *Dad* (*Daughter against Dpp*), encoding the *Drosophila* homolog of Smad6/7 (Fig. 3A). Thus to test whether *Dad* is the specific gene that accounts for the modification of the CDK8-specific phenotypes by these two *Df* lines, we performed similar genetic tests with a mutant allele of *Dad*, *Dad^MI04922^*. Indeed, *Dad^MI04922^* dominantly enhanced the CDK8-RNAi (Fig. 3B), CycC-RNAi (Fig. 3C), or CDK8-RNAi plus CycC-RNAi (Fig. 3D) phenotypes, but suppressed the CDK8-overexpression phenotype (Fig. 3E). These effects on the CDK8-specific vein phenotypes are similar to those observed for *Df(3R)BSC748* and *Df(3R)Exel6176*, suggesting that *Dad* is the specific gene that genetically interacts with CDK8 *in vivo*.

**Fig. 3.**
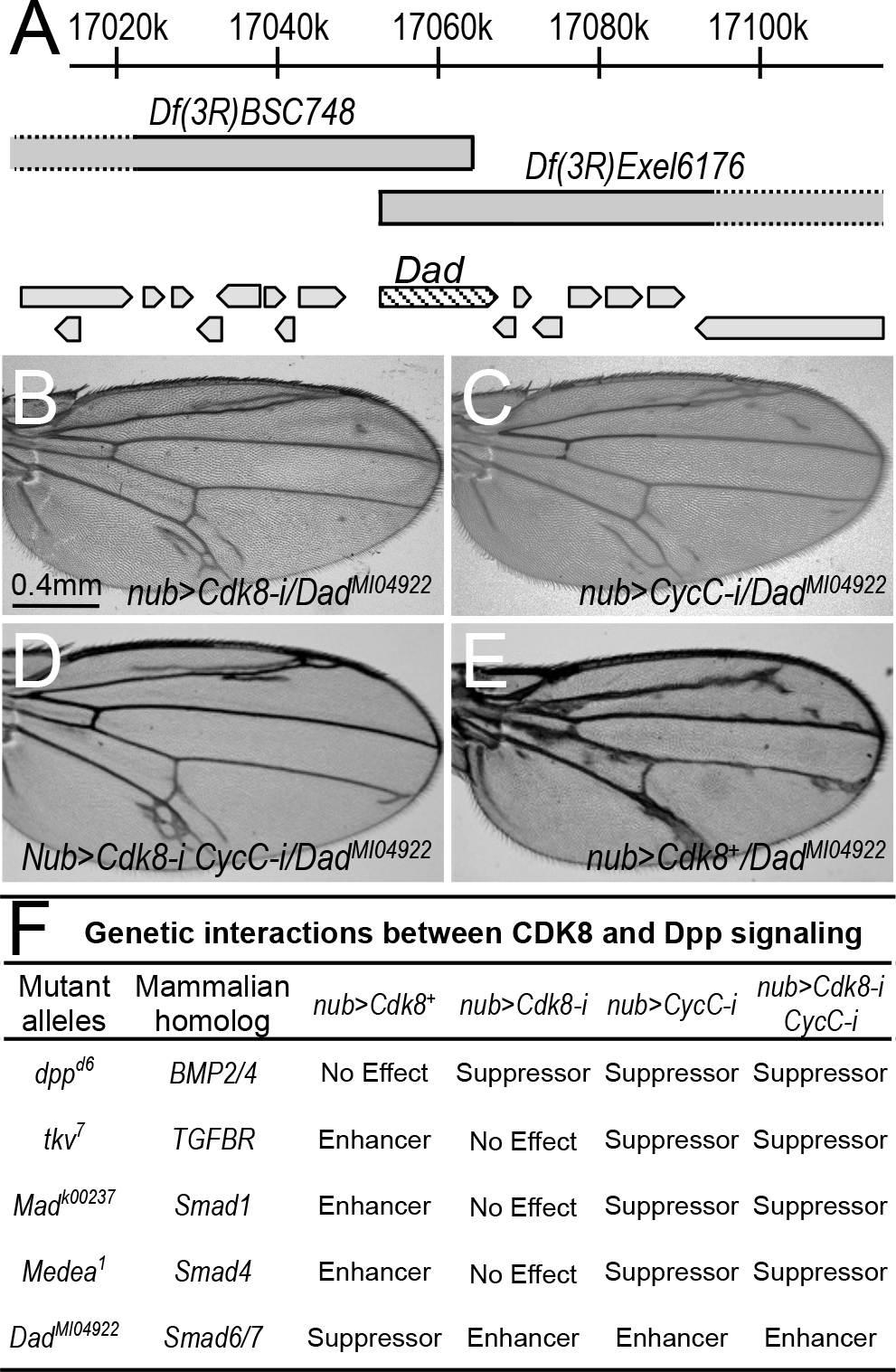
Identification of Dad as an enhancer of the *nub>Cdk8i* and *nub>CycCi* phenotypes but a suppressor of Cdk8-overexpression phenotype. (A) Genome region schematic diagram of *Df(3R)BSC748* and *Df(3R)Exel6176* uncovering the same gene, *dad*. Adult female wings of (B) *nub-Gal4/+; UAS-Cdk8-RNAi/ Dad^MI04922^*; (C) *nub-Gal4/+; UAS-CycC-RNAi/Dad^MI04922^*; (D) *nub-Gal4/+; UAS-Cdk8-RNAi CycC-RNAi /Dad^MI04922^*; and (E) *nub-Gal4>UAS-Cdk8^+^/+; Dad^MI04922^/+*. Scale bar in B: 0.4mm. (F) Summary of genetic interactions between CDK8-CycC and components of the Dpp signaling pathway.

### Mutants of multiple components of the Dpp signaling pathway genetically interact with CDK8-CycC

The protein Dad functions as an inhibitory Smad in the Dpp/TGFβ signaling pathway, which plays critical roles in regulating cell proliferation and differentiation during the development of metazoans (35-40). During the development of the wing discs, Dpp spreads from the anterior-posterior boundary to the anterior half and posterior halves (35-37). Upon binding of the Dpp ligand to the Tkv-Punt receptor complex on the cell membrane, the TGFβ type II receptor Punt phosphorylates and activates the type I receptor Tkv. This results in the phosphorylation of Mad by Tkv at its C-terminal SSXS motif, known as the phospho-Mad protein or pMad. Medea, the unique co-Smad protein in *Drosophila*, associates with pMad in the cytoplasm, and then this heteromeric Smad complex translocates into the nucleus and regulates the expression of its target genes (39, 41-43).

The genetic interactions between CDK8-CycC and Dad prompted us to test whether mutant alleles of other components of the Dpp signaling pathway could also genetically interact with CDK8 and CycC. For this, we crossed multiple mutant alleles of these components with the CDK8-CycC depletion or overexpression lines. As summarized in Fig. 3F, mutants of multiple components of the Dpp signaling pathway could either dominantly enhance or suppress the CDK8-specific vein phenotypes. For example, *dpp^d6^, tkv^7^, Mad^k00237^*, and *Medea^1^* all dominantly suppressed the ectopic vein phenotype caused by depletion of CycC, or both CDK8 and CycC; mutant of *dpp^d6^* could also suppress the CDK8-RNAi phenotype; however, *tkv^7^, Mad^k00237^*, and *Medea^1^* enhance the CDK8-overexpression phenotype (Fig. 3F). Taken together, these genetic interactions suggest that CDK8-CycC may affect vein patterning by modulating Dpp signaling.

### CDK8-CycC positively regulates Mad-dependent transcription

Given that CDK8 and CycC are known subunits of the Mediator complex, which serves as a scaffold complex mediating the interactions between the RNA Pol II basal transcription initiation apparatus and a variety of gene-specific transcription activators (3, 7, 44), the most parsimonious model to explain the genetic interactions between Dpp signaling and CDK8-CycC is that the CDK8-CycC complex may directly regulate the transcriptional activity of Mad in the nucleus. To test this model, we analyzed the effects of CDK8-CycC depletion on the expression of *spalt* (*sal*), a well-characterized direct target gene of Mad involved in vein differentiation (45-47). Specifically, the expression of *sal-lacZ* serves as a reporter for the transcriptional activity of Mad, downstream of the Dpp signaling pathway (48).

Because the expression of *sal-lacZ* is symmetric along the dorsal-ventral boundary of the wing pouch area of the wing discs (Fig. 4A), we tested whether specific depletion of CDK8 or CycC within the dorsal compartment of the wing discs could affect the transcriptional activity of Mad by detecting the transcription level of *sal* using an anti-β-galactosidase (anti-β-Gal) antibody. For this, we depleted CDK8, CycC, or both, using the *ap-Gal4* driver, and then compared the β-Gal expression between the dorsal and ventral compartments. Depletion of CDK8 (Fig. 4B), CycC (Fig. 4C), or both (Fig. 4D), the dorsal compartment significantly decreased β-Gal expression level at the dorsal compartment compared with the ventral compartment of the same disc. As expected, depleting Dpp (Fig. S2A), Mad (Fig. S2B), or Medea (Fig. S2C) using the same approach reduced the expression of the *sal-lacZ* reporter in the dorsal compartment. These observations suggest that the CDK8-CycC complex positively regulates Mad-dependent transcription.

**Fig. 4.**
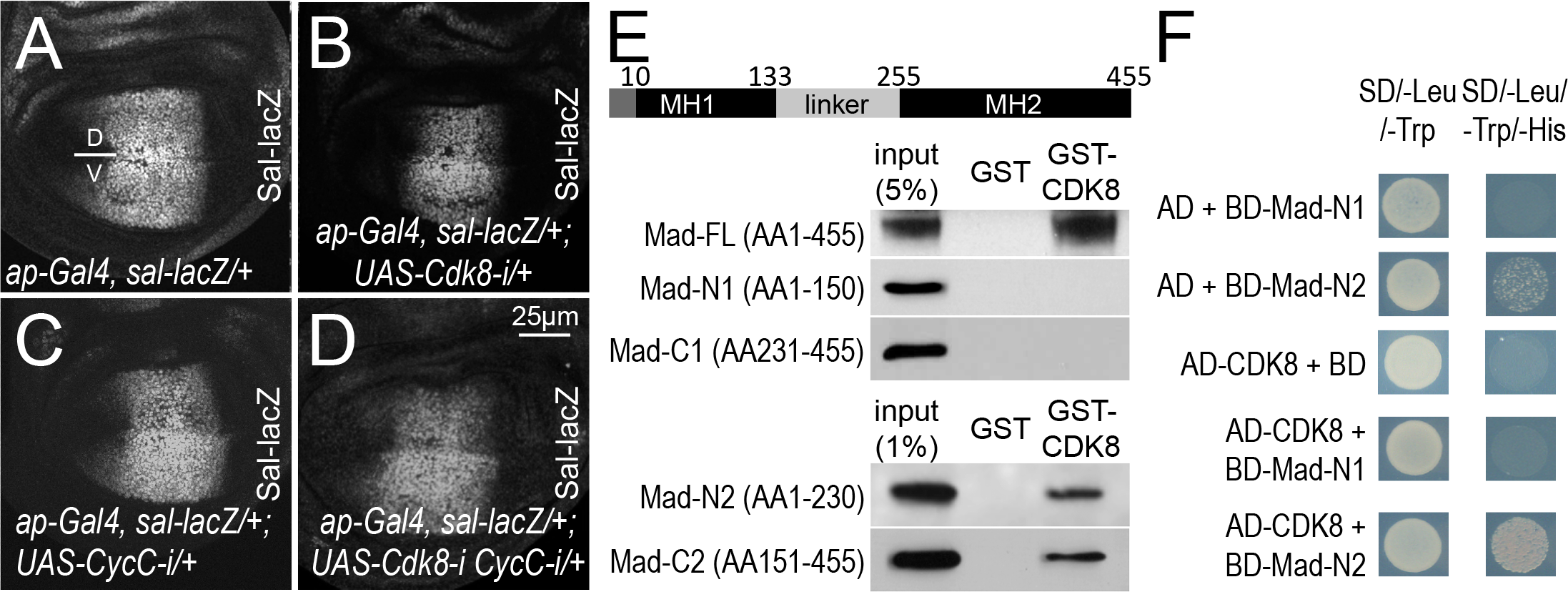
CDK8-CycC may positively regulate Mad/Smad-dependent transcription. Confocal images of the wing pouch area of a L3 wondering larvae wing disc of (A) *ap-Gal4, sal-lacZ/+* (control); (B) *ap-Gal4, sal-lacZ/+; UAS-Cdk8-i/+*; (C) *ap-Gal4, sal-lacZ/+; UAS-CycC-i/+*; and (D) *ap-Gal4, sal-lacZ/+; UAS-Cdk8-i CycC-i/+*. All signals presented were from anti-βgalactosidase staining. Scale bar in D: 25µm. (E) Western Blots of a GST pulldown assay between GST-CDK8 and His-tagged Mad fragments. The amino acids positions of MH1 (Mad homology domain 1) and MH2 (Mad homology domain 2), separated by the linker region, are based on BLAST search of *Drosophila* Mad-RA isoform (455AA). The other isoform, Mad-RB (525AA), has additional 70AA at the N-terminus. We focused on the Mad-RA isoform in this work. (F) Yeast two-hybrid assay showing the specific interaction between CDK8 and the linker region of Mad. SD/-Leu/-Trp and SD/-Leu/-Trp/-His are dropout media lacking “Leu and Trp”, or “Leu, Trp, and His”, respectively. The con-transformed yeast cultures were spotted on SD/Leu/-Trp and SD/-Leu/-Trp/-His plates, positive interactions results in yeast growth on the SD/-Leu/-Trp/-His plate. AD, GAL4-activation domain (prey); BD, GAL4-DNA-binding domain (bait); AD- or BD-protein, AD- or BD-fusion proteins.

Since Mad phosphorylation at its C-terminus (pMad) by the Tkv-Punt receptor complex marks the activation of Mad (Fig. 4E), we tested whether CDK8 affects the pMad level. For this, we depleted CDK8, CycC, or both, with the *ap-Gal4* line, and then detected the levels of the activated Mad with an anti-pMad antibody. In the wing pouch area of the control discs, the pMad protein is symmetrically distributed along the dorsal-ventral boundary (Fig. S3A). However, depletion of CDK8-CycC did not affect pMad levels when comparing the dorsal compartment with the ventral compartment (Figs. S3B’-S3D’), suggesting that CDK8-CycC does not affect the phosphorylation of Mad at its carboxy terminus in the cytoplasm. These results support the idea that the CDK8-CycC complex may directly regulate the transcriptional activity of Mad in the nucleus.

### Direct interactions between CDK8 and Mad

R-Smads are characterized with a highly conserved amino-terminal MH1 (Mad homology 1) domain that binds to DNA, a C-terminal MH2 (Mad homology 2) domain that harbors the transactivation activity, separated by a serine- and proline-rich linker region (Fig. 4E) (49). It was previously reported that CDK8 and a few other kinases (see below) may directly phosphorylate Smad proteins both in *Drosophila* and mammalian cells (14, 15, 30, 42, 49), but whether and how CDK8 interacts with Smads remains unknown. To determine whether CDK8 directly interacts with Mad, we performed a GST-pulldown assay. As shown in Fig. 4E, purified GST-CDK8 can directly bind with His-tagged full length Mad (Mad-FL, AA1-455) expressed in *E. coli*. We then further mapped the specific domain of Mad that interacts with CDK8 using His-tagged fragments of the Mad protein (see Materials and Methods for details). We have observed that the “Mad-N2” fragment (AA1-230) and the “Mad-C2” fragment (AA151-455), but not the “Mad-N1” fragment (AA1-150) or the “Mad-C1” fragment (AA231-455), can directly interact with CDK8 (Fig. 4E). We validated the interaction between CDK8 and the linker region using the yeast two-hybrid (Y2H) assay: the “Mad-N2” fragment, but not the “Mad-N1” fragment, as the bait can interact with full length CDK8 as the prey (Fig. 4F). It is not feasible to use this two-hybrid approach test Mad-FL or Mad-C1/C2 fragments as baits, since they auto-activate as the baits; while using full-length CDK8 as the bait is also able to auto-activate (Fig. S4). Taken together, these results suggest that CDK8 interacts directly with part of the linker region of Mad protein (AA151-230). Implications of these physical interactions are further discussed below.

### Involvement of additional subunits of the Mediator complex in regulating the Mad/Smad-dependent transcription

Interestingly, Med15/ARC105 subunit of the Mediator complex directly interacts with the transactivation MH2 domain of Smad2/3, thereby mediating the Smad2/3-Smad4-dependent transcription in *Xenopus* (50). Med15 has been previously shown to be required for the transcription of Dpp target genes in *Drosophila (51)*. However, whether other Mediator subunits are involved in regulating the Mad/Smad-dependent transcription remains unknown. To address this question, we depleted individual subunits of the Mediator complex with *ap-Gal4* and then analyzed the expression of the *sal-lacZ* reporter. Of the 30 Mediator subunits tested (Fig. 5A), we have observed that depletion of six additional Mediator subunits by *ap-Gal4*, including Med12 (Fig. 5B), Med13 (Fig. 5C), Med15 (Fig. 5D), Med23 (Fig. 5E), Med24 (Fig. 5F), and Med31 (Fig. 5G), significantly reduced the expression of *sal-lacZ* in the dorsal cells compared with the cells in the ventral compartment of the same wing discs, similar to depletion of CDK8 or CycC. These results suggest that these Mediator subunits are required for the Mad-activated gene expression. However, RNAi depletion of the remaining 15 Mediator subunits did not significantly affect *sal-lacZ* expression (Fig. 5A), since β-Gal expression remained symmetric along the dorsal-ventral boundary as exemplified for Med1 (Fig. 5H) and Med25 (Fig. 5I). Furthermore, depleting the remaining Mediator subunits, including Med7 (Fig. 5J), Med8 (Fig. S5A), Med14 (Fig. S5B), Med16 (Fig. S5C), Med17 (Fig. S5D), Med21 (Fig. S5E), and Med22 (Fig. S5F) strongly disrupted the morphology of the wing discs, making it difficult to determine whether these subunits affect *sal* transcription. Taken together, these observations suggest that multiple Mediator subunits, but apparently not all of them, are required for Mad-dependent transcription in *Drosophila*.

**Fig. 5.**
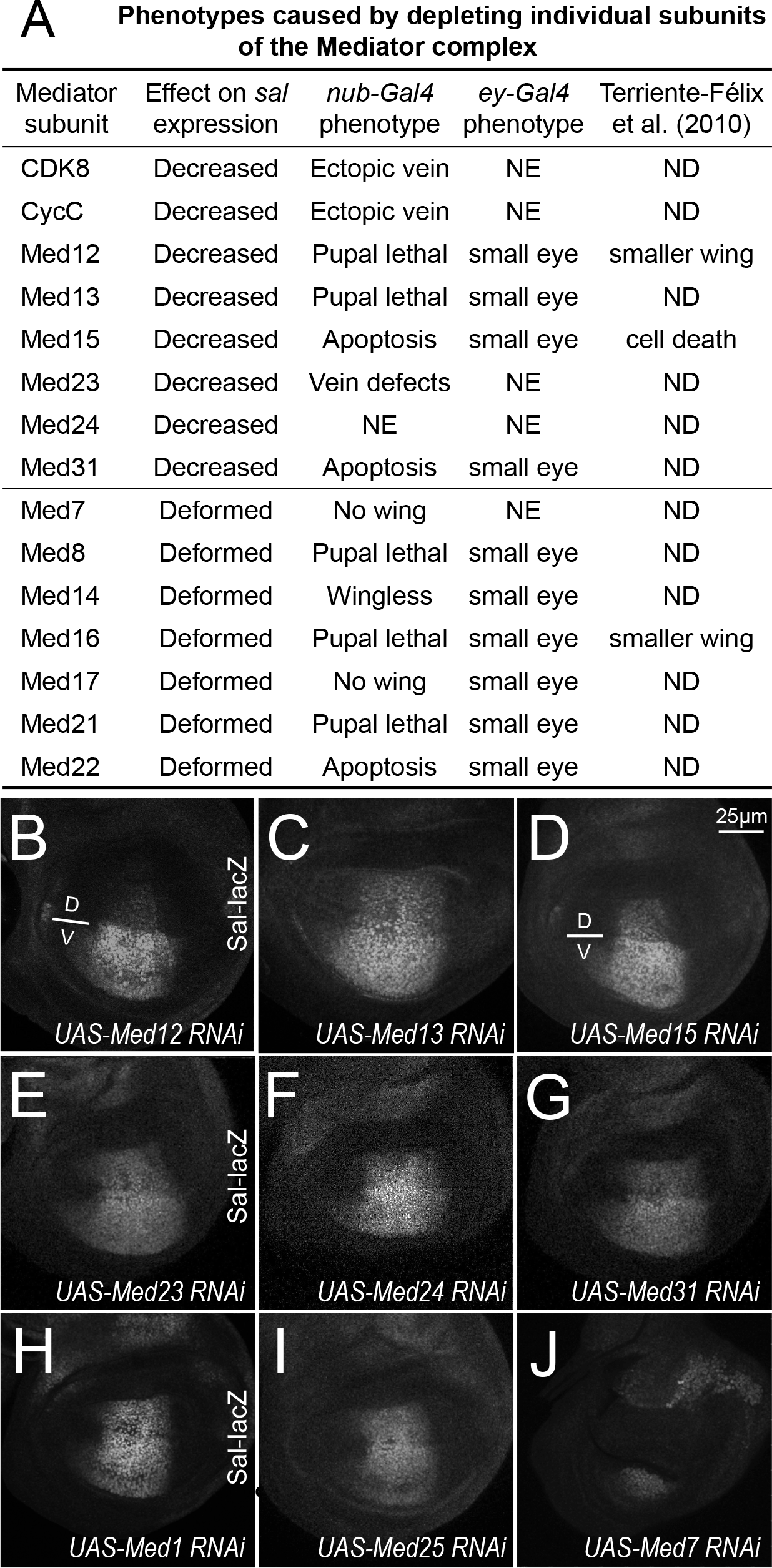
Involvement of additional Mediator subunits in regulating Mad/Smad-dependent transcription. (A) Summary of effects on *sal* transcription and morphology of wing and eye by tissue-specific depleting individual subunits of the Mediator complex: depleting eight of them decreased *sal* expression, while depleting seven of them disrupted the morphology of wing discs. NE: no effects; ND: not determined. Confocal images of anti-β-Gal staining of wing discs of the following genotypes: (B) *ap-Gal4, sal-lacZ/+; UAS-Med12 RNAi/+*; (C) *ap-Gal4, sal-lacZ/+; UAS-Med13 RNAi/+*; (D) *ap-Gal4, sal-lacZ/+; UAS-Med15 RNAi/+*; (E) *ap-Gal4, sal-lacZ/+; UAS-Med23 RNAi/+*; (F) *ap-Gal4, sal-lacZ/+; UAS-Med24 RNAi/+*; (G) in *ap-Gal4, sal-lacZ/+; UAS-Med31 RNAi/+*; (H) *ap-Gal4, sal-lacZ/+; UAS-Med1 RNAi/+*; (I) *ap-Gal4, sal-lacZ/UASMed25 RNAi*; and (J) *ap-Gal4, sal-lacZ/+; UAS-Med7 RNAi/+*. Scale bar in D: 25µm.

### CDK9 and Yorkie also positively regulate the Mad/Smad-dependent transcription

Besides CDK8, several other kinases, such as CDK7, CDK9, GSK3 (Glycogen synthase kinase 3), and MAPKs (mitogen-activated protein kinases) such as ERK and ERK2 (extracellular signal-regulated kinases), have been implicated to phosphorylate and regulate the transcriptional activity of Smads (14, 15, 49, 52). The four phosphorylation sites (Ser/Thr residues) within the linker region of Smads appear to be conserved from *Drosophila* to mammals (Fig. 6A). The phosphorylation of Smads within the linker region may facilitate the subsequent binding with transcription co-factors, such as YAP (Yes-associated protein) (14). However, it is still unclear whether all of these kinases regulate the Smads activity *in vivo*, and with the exception of YAP (Yorkie or Yki, in *Drosophila*), it is also unclear whether these regulatory mechanisms are conserved during evolution.

**Fig. 6.**
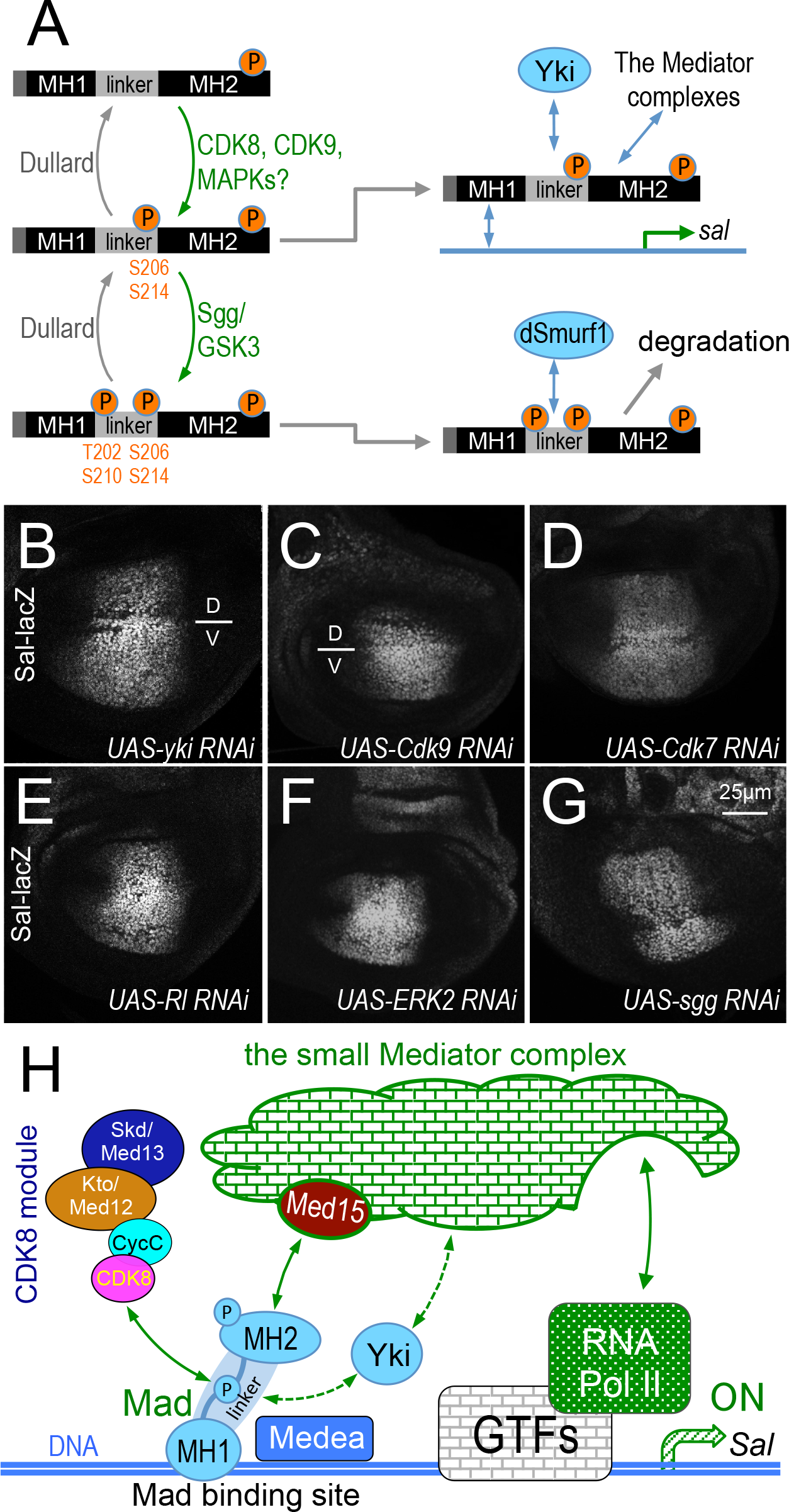
CDK9 and Yorkie also positively regulate Mad/Smad-dependent transcription. (A) Model: linker region of pMad may be phosphorylated by CDK8, CDK9, or MAPKs as priming kinase recruiting Yki/YAP binding to pMad to drive target gene, such as *sal* transcription; and further phosphorylation by Sgg/GSK3 at the linker region may switch the binding to dSmuf1 and causes pMad degradation. Anti-β-Gal staining of wing discs of the following genotypes: (B) *ap-Gal4, sal-lacZ/+; UAS-yki RNAi/+*; (C) *ap-Gal4, sal-lacZ/UAS-Cdk9 RNAi*; (D) *ap-Gal4, sallacZ/UAS-Cdk7 RNAi*; (E) *ap-Gal4, sal-lacZ/+; UAS-rl RNAi/+*; (F) *ap-Gal4, sal-lacZ/+; UASERK2 RNAi/+*; and (G) *ap-Gal4, sal-lacZ/UAS-sgg RNAi*. Scale bar in G: 25µm. (I) Model of Mad/Smad-dependent transcription activation through the CDK8 module and the Mediator complex. (GTFs: General Transcription Factors)

To validate the relevance of these kinases in regulating Mad-dependent gene expression, we depleted the *Drosophila* orthologs of CDK7, CDK9, Shaggy (Sgg, the GSK3 homolog in *Drosophila*), Rolled and dERK2 (MAPK/ERK homologs in *Drosophila*), in the dorsal compartment of wing discs, and then analyzed the expression of *sal-lacZ* in the wing pouch. For this analysis, we used Yki as a positive control (14). Depletion of either Yki (Fig. 6B) or CDK9 (Fig. 6C) in the dorsal cells significantly reduced the expression of *sal-lacZ* compared to the cells in the ventral compartment of the same discs. These observations suggest that both Yki and CDK9 are required for Mad/Smad-dependent transcription in *Drosophila*, which is consistent to the previous reports (14, 30). However, depletion of CDK7 (Fig. 6D) or *Drosophila* MAPK homologs, either Rolled (Fig. 6E) or dERK2 (Fig. 6F), did not affect the expression of *sal-lacZ*. Although depletion of Sgg increased the size of the dorsal compartment, the intensity of anti-β-Gal staining remained similar to the ventral compartment (Fig. 6G). Together with the previous reports (14, 49), our *in vivo* analyses suggest that the roles of CDK8, CDK9, and Yki/YAP on the Mad/Smad-dependent transcription are conserved between mammals and *Drosophila*.

## Discussion

To study the function and regulation of CDK8 *in vivo*, we have developed a genetic system that yields robust readouts for the CDK8-specific activities in developing *Drosophila* wings. These genetic tools provide a unique opportunity to perform a dominant modifier genetic screen, which allow us to identify multiple components of the Dpp signaling pathway that can genetically interact with CDK8 and CycC *in vivo*. Our subsequent genetic and cellular analyses reveal that CDK8, CycC, and six additional subunits of the Mediator complex, as well as CDK9 and Yki are required for the Mad-dependent transcription in the wing imaginal discs. In addition, CDK8 can directly interact with the linker region of Mad. These results have extended the previous biochemical and molecular analyses on how different kinases and transcription cofactors modulate the Mad/Smad-activated gene expression in the nucleus. Further mapping of specific genes uncovered by other deficiency lines may also open up the new directions to advance our understanding about the conserved functions of CDK8-CycC during development.

### Multiple subunits of the Mediator complex are required for Mad/Smad-dependent transcription

The Mediator complexes function as scaffolds physically bridging gene-specific transcription factors to the RNA Pol II general transcription apparatus, and diverse transactivators have been shown to interact directly with distinct Mediator subunits (4, 6-9, 53). However, it is unclear whether all Mediator subunits are required by different transactivators to regulate gene expression, or whether Mediator complexes composed of fewer and different combinations of Mediator subunits exist in differentiated tissues or developmental stages. Gene-specific combinations of the Mediator subunits may be required in different transcription processes, as not all Mediator subunits are simultaneously required for all transactivation process (54). For example, ELK1 target gene transcription requires Med23, but lacking Med23 does not functionally affect some other ETS transcription factors, e.g. Ets1 and Ets2 (55). Similarly, Med15 is required for the expression of Dpp target genes, but does not appear to affect the expression of EGFR and Wg targets in *Drosophila* (51).

It has been previously reported that the Med15 subunit is required for the Smad2/3-Smad4 dependent transcription, as its removal from the Mediator complex abolishes the expression of Smad-target genes and disrupts Smad2/3-regulated dorsal-ventral axis formation in *Xenopus* embryos (50). Further biochemical analyses showed that increased Med15 enhances, while its depletion decreases, the transcription of Smad2/3 target genes, and that the Med15 subunit can directly bind to the MH2 domain of Smad2 or Smad3 (50). In *Drosophila*, loss or reduction of Med15 reduced the expression of Dpp targets, resulting in smaller wings and disrupted vein patterning (mainly L2) (51). We also observed that depletion of Med15 or CDK8 reduces the expression of a Mad-target gene. These observations support the idea that CDK8 and Med15 play a conserved and positive role in regulating Mad/Smad-activated gene expression.

Aside from Med15 and CDK8, it remained unclear whether other Mediator subunits are also involved in Mad/Smad-dependent transcription. We identified six additional Mediator subunits that are required for the Mad-dependent transcription, including CycC, Med12, Med13, Med23, Med24, and Med31 (Fig. 5A). Interestingly, except Med15, counterparts of the other seven subunits are not essential for cell viability in the budding yeast (5). The similar effects of the four CKM subunits on Mad-activity suggest that they may function together to stimulate the Mad-dependent transcription. We note that depletion of seven Mediator subunits, e.g. Med7, Med8, Med14, Med16, Med17, Med21, and Med22, severely disrupted the morphology of the wing discs (Fig. 5A), making it difficult to assay their effects on the transcriptional activity of Mad *in vivo*. Except Med16, all corresponding subunits are critical for cell viability in the budding yeast (5). In contrast, reducing expression of the 15 remaining subunits of the *Drosophila* Mediator complex did not significantly alter the expression of a Mad-dependent reporter (Table S3). Med1 and Med25 are loosely associated to the small Mediator complex in human cell lines (5). A caveat for these negative results is that the RNAi lines may not be effective enough to affect *sal-lacZ* expression, even though the transgenic RNAi lines for majority of these subunits cam generate phenotypes in the eye, wing or both (Table S3). Therefore, these results indicate that not all Mediator subunits are required for the Mad-dependent gene expression in the developing wing discs.

### Role of Yki/YAP and different kinases in regulating Mad/Smad-dependent transcription

Interestingly, Yki/YAP, which can function as a transcriptional co-factor for Mad/Smad, was also reported to associate with several subunits of the Mediator complex to drive transcription. For example, Med12, Med14, Med23, and Med24 were identified from a YAP IP-mass spectrometry sample from HuCCT1 cells (56). Med23 was also reported to regulate Yki-dependent transcription of *Diap1* in the *Drosophila* wing discs (57). In this work, we found that Yki, Med12, Med23, and Med24 were also required for the Mad-dependent transcription of *sallacZ*. Although the exact molecular mechanisms of how Yki interacts with certain Mediator subunits remain unclear, it is plausible that Yki may further strengthen the binding between Mad and Med15 through interactions with other subunits such as Med12, Med23, and Med24.

Based on biochemical analyses of the Smad1 phosphomutants and cell biological analyses using cultured human epidermal keratinocytes (HaCaT cells), several kinases including CDK8, CDK9, and ERK2 were shown to phosphorylate serine residues (S) within the linker region of pSmad1 at S186, S195, S206, and S214, or the equivalent sites in pSmad2/3/5. These modifications were proposed to positively regulate the Smad1-dependent transcriptional activity (14). Of these sites, S206 and S214 are both conserved from *Drosophila* to humans (Fig. S6). In addition, studies using *Xenopus* embryos and cultured L cells suggest that MAPKs may phosphorylate the linker region of Smad1 (including S214) and lead to its degradation (52). Nevertheless, analyses with *Drosophila* embryos or wing discs indicate that S212 (equivalent to human pSmad1 S214) is phosphorylated by CDK8, and S204 (unique in *Drosophila*) or S208 (equivalent to human pSmad1 S210) are phosphorylated by Sgg/GSK3 (15). These studies suggest the following model to explain how Smads activate the expression of their target genes and how this process is turned off (Figs. 6A, 6H): after Smads are phosphorylated at their C-termini and translocate into the nucleus, CDK8 and CDK9 (potentially alsoMAPKs) act as the priming kinases to further phosphorylate pSmads in the linker region at S206 and S214, which may facilitate the interaction between pSmads and transcription cofactors such as YAP, thereby stimulating the expression of Smads target genes. Subsequently, pSmads are further phosphorylated by GSK3 within the linker region at T202 and S210, which may facilitate Smad1/5 binding to E3 ligases such as Smurf1 and Nedd4L, thereby causing the degradation of Smads through the ubiquitin-proteasome pathway (14, 15, 30, 42, 49).

This model explains how the transactivation of Smads is coupled to its degradation, similar to other transcriptional activators (58). However, it is rather challenging to determine whether these kinases act redundantly or specifically for different phosphorylation sites, the exact orders of these phosphorylation events, as well as their biological consequences *in vivo*. Moreover, it remains unexplored whether these regulatory mechanisms are conserved during evolution. The importance of these issues is highlighted by the critical roles of TGFβ signaling in regulating the normal development of metazoans and the dysregulation of this pathway in a wide variety of human diseases such as cancers (40, 59, 60).

The precise spatiotemporal activation of the Dpp signaling pathway in the wings discs is critical for proper formation of the stereotypical vein patterns in *Drosophila* (45). This model system provides an ideal opportunity to dissect the dynamic regulation of the Mad-activated gene expression in the nucleus. Indeed, depleting CDK8 in wing discs reduces the expression of the Mad-dependent *sal-lacZ* reporter, suggesting that CDK8 positively regulates Mad-dependent transcription, which is consistent to the effects of CDK8 on Smad1/5-dependent transcription in mammals (14, 61). Depleting CDK8 does not affect the phosphorylation of Mad at its C-terminus as revealed by pMad immunostaining (Fig. S3), as well as the physical interaction between CDK8 and the linker region of Mad, supporting the idea that CDK8 may only affect the subsequent phosphorylation of Mad.

Besides CDK8, depleting CDK9 also decreased the expression of the *sal-lacZ* reporter, supporting the notion that CDK8 and CDK9 may play non-redundant roles in further phosphorylating pMad in the nucleus. However, we did not observe any effects of depletion of MAPKs on *sal*-*lacZ* expression, suggesting that their role in regulating the transcriptional activity of Smads may not be conserved in *Drosophila*. Alternatively, the two MAPK/ERK homologs, Rolled and ERK2, may act redundantly in regulating Mad-dependent transcription. Lastly, depleting Sgg/GSK3 in the dorsal compartment of the wing disc increases the size of this compartment, yet the expression level of the *sal-lacZ* reporter is similar to the ventral compartment. These observations are consistent to the previously report that phosphorylation of Mad in the linker regions by CDK8 and Sgg/GSK3 regulate the level and range of Mad-dependent gene expression (14, 15, 30, 42, 49).

Together with the previous reports (14, 15, 30, 42, 49, 62), our data support that CDK8 or CDK9 may phosphorylate pMad at the linker region, which may facilitate the binding between Yki and Mad. We speculate that this interaction may synergize the recruitment of the Mediator complex, presumably through at least the interaction between its Med15 subunit and the MH2 domain of Mad. Alternatively, Yki may also facilitate the recruitment of the whole Mediator complex through its interactions with Med12, Med23, and Med24. The synergistic interactions among Mad, Yki, the Mediator complex, and RNA Pol II may be required for the optimal transcriptional activation of the Mad-target genes (Fig. 6H).

One of the important future challenges is to illuminate the dynamic interactions between these factors and diverse protein complexes that couple the transactivation effects of Mad/Smads on gene transcription with their subsequent degradation at the molecular level. Smad3 phosphorylation strongly correlates with Med15 levels in breast and lung cancer tissues, together they potentiate metastasis of breast cancer cells (63). Thus it will be important to test whether similar effects of the additional Mediator subunits that we identified here can be observed in mammalian cells. It will also be interesting to determine whether a partial Mediator complex, composed of a few Mediator subunits, exists and regulates Mad/Smad-dependent gene expression in the future. Furthermore, detailed biochemical analyses may yield mechanistic insights into how CDK8 and Med15 act in concert in stimulating the Mad/Smad-dependent gene expression.

### Identification of novel genomic loci that genetically interact with CDK8 *in vivo*

To understand how dysregulated CDK8 contributes to a variety of human cancers, it is essential to elucidate the function and regulation of CDK8 *in vivo*. Given that CDK8-CycC and other subunits of the Mediator complex are conserved in almost all eukaryotes (5), *Drosophila* serves as an ideal model system to identify both the upstream regulators and the downstream effectors of CDK8 *in vivo*. Our dominant modifier genetic screen is based on the vein phenotypes caused by specific alteration of CDK8 activity in the developing wing disc, which serves as a unique *in vivo* readout for the CDK8-specific activities in metazoans. This screen led us to identify 26 genomic regions that include genes whose haplo-insufficiency can consistently modify CDK8-CycC depletion or CDK8-overexpression phenotypes. Identification of *Dad* and genes encoding additional components of the Dpp signaling pathway provides a proof of principle for this approach. Since each of the chromosomal deficiencies uncovers multiple genes, further mapping of the relevant genome regions is expected to identify the specific genetic loci encoding factors that may function either upstream or downstream of CDK8 *in vivo*. Further analyses of the underlying molecular mechanisms in both *Drosophila* and mammalian systems will advance our understanding of how dysregulation of CDK8 contributes to human diseases, and may also aid the development of novel therapeutic approaches.

## Materials and Methods

### Fly strains

Flies were raised on standard cornmeal, molasses and yeast medium, and all genetic crosses were maintained at 25°C. The *UAS-Cdk8^+^* and *UAS-Cdk8^KD^* lines were generated using the pUASt vector (32). The construct allowing conditional expression of a kinase-dead CDK8 form (D173A; (64)) was generated through site-specific mutagenesis by double PCR, using the overlap extension method. The *UAS-Cdk8-RNAi*, *UAS-CycC-RNAi*, and *UAS-Cdk8-RNAi CycC-RNAi* line was generated using the pVALIUM20 vector (65).

We obtained the following strains from the Bloomington *Drosophila* Stock Center: *nub-Gal4* (BL-25754), *ap-Gal4* (BL-3041), *UAS-CDK7-RNAi* (BL-57245), *UAS-Cdk8-RNAi* (BL-67010), *UAS-CDK9-RNAi* (BL-34982), *UAS-CycC-RNAi* (BL-33753), *UAS-dpp-RNAi* (BL-33618), *UAS-2xEGFP* (BL-6874), *UAS-erk-RNAi* (BL-34744), *UAS-mad-RNAi* (BL-43183), *UAS-medea-RNAi* (BL-43961), *UAS-rl-RNAi* (BL-34855), *sal-lacZ* (BL-11340), *UAS-sgg-RNAi* (BL-38293), and *UAS-yki-RNAi* (BL-34067). In addition, we tested the following mutant alleles of the Dpp signaling pathway: *dpp^d6^/CyO* (BL-2062), *tkv^7^/CyO* (BL-3242), *Mad^k00237^/CyO* (BL-14578), *Medea^1^/TM3* (BL-9033), and *Dad^MI04922^/TM3* (BL-37913).

The following RNAi stocks, generated by the *Drosophila* TRiP project (65), were used to deplete the subunits of the Mediator complex: *UAS-Med1-RNAi* (BL-34662), *UAS-Med4-RNAi* (BL-34697), *UAS-Med6-RNAi* (BL-33743), *UAS-Med7-RNAi* (BL-34663), *UAS-Med8-RNAi* (BL-34926), *UAS-Med9-RNAi* (BL-33678), *UAS-Med10-RNAi* (BL-34031), *UAS-Med11-RNAi* (BL-34083), *UAS-Med12-RNAi* (BL-34588), *UAS-Med13-RNAi* (BL-34630), *UAS-Med14-RNAi* (BL-34575), *UAS-Med15-RNAi* (BL-32517), *UAS-Med16-RNAi* (BL-34012), *UAS-Med17-RNAi* (BL-34664), *UAS-Med18-RNAi* (BL-42634), *UAS-Med19-RNAi* (BL-33710), *UAS-Med20-RNAi* (BL-34577), *UAS-Med21-RNAi* (BL-34731), *UAS-Med22-RNAi* (BL-34573), *UAS-Med23-RNAi* (BL-34658), *UAS-Med24-RNAi* (BL-33755), *UAS-Med25-RNAi* (BL-42501), *UAS-Med26-RNAi* (BL-28572), *UAS-Med27-RNAi* (BL-34576), *UAS-Med28-RNAi* (BL-32459), *UAS-Med29-RNAi* (BL-57259), *UAS-Med30-RNAi* (BL-36711), and *UAS-Med31-RNAi* (BL-34574).

To facilitate the dominant modifier genetic screen and the subsequent analyses, we generated the following strains using the standard *Drosophila* genetics: “*w^1118^; nub-Gal4>UAS-Cdk8^+^/CyO*” (i.e., “*nub>Cdk8^+^/CyO*” line), “*w^1118^; nub-Gal4; UAS-Cdk8-RNAi*” (i.e., “*nub>Cdk8-i*” line), “*w^1118^; nub-Gal4; UAS-CycC-RNAi*” (i.e., “*nub>CycC-i*” line), “*w^1118^; nub-Gal4; UAS-Cdk8-RNAi CycC-RNAi*” (i.e., “*nub>Cdk8-i CycC-i*” line), and “*w^1118^; ap-Gal4, sallacZ/T(2:3)*”. All deficiency (*Df*) lines, listed in Table S1 and Table S2, were obtained from the Bloomington *Drosophila* stock center.

For the *Df* lines in the X chromosome, we crossed *Df* female virgins with males of with the “*nub>Cdk8^+^/CyO*”, “*nub>Cdk8-i*”, “*nub>CycC-i*”, or “*nub>Cdk8-i CycC-i”* stocks. For the *Df* lines in the second and third chromosomes, the *Df* males were crossed with female virgins of the aforedescribed stocks carrying the CDK8-specific phenotypes. The control crosses were performed using *w^1118^* males or female virgins. For all these crosses, the wing vein patterns in all F1 females without any balancer chromosomes were inspected under dissecting microscopes for potential dominant modifications. For example, we crossed *Df(1)BSC531, w^1118^/FM7h* female virgins with “*w^1118^/Y; nub>Cdk8^+^/CyO*” males, and then scored F1 females with the following genotype: “*Df(1)BSC531, w^1118^/ w^1118^; nub>Cdk8^+^/+*”. Similarly, we crossed “*w^1118^; nub-Gal4; UAS-Cdk8-RNAi*” female virgins with “*Df(2R)Exel6064/CyO*” males, and then scored F1 females with the following genotype: “*w^1118^/+; nub-Gal4/Df(2R)Exel6064; UAS-Cdk8-RNAi/+*”.

### Adult *Drosophila* wing imaging

The wings from adult females were dissected onto slides, briefly washed using isopropanol, and then mounted in 50% Canada balsam in isopropanol. Images were taken under 5X objective of a microscope (Leica DM2500) and then processed by Adobe Photoshop CS6 software.

### Immunocytochemistry

Wing discs from third instar larvae at the late wandering stage were dissected and fixed in 5% formaldehyde at room temperature for 30 minutes. After rinsing with PBS-Triton X-100 (0.2%), the samples were blocked in PBS-Triton X-100-NGS-BSA (PBS+0.2% Triton X-100+5% Normal Goat Serum+0.2% Bovine Serum Albumin) at room temperature for one hour. For immunostaining of CDK8 and CycC, we used an anti-dCDK8 (1:2000) antibody and an anti-dCycC (1:2000), both generated in guinea pigs (66), and diluted in PBS-Triton X-100-NGS-BSA. The expression of the *lacZ* reporter expression was detected using an anti-β-galactosidase monoclonal antibody (1:50 in PBS-Triton X-100-NGS-BSA; obtained from the Developmental Studies Hybridoma Bank, DSHB-40-1a-s). C-terminal phosphorylated Mad (equivalent sites to hSmad3 S423+S425) was detected by anti-pSmad3 (1:500 in PBSTriton X-100-NGS-BSA; purchased from Abcam, ab118825). Wing discs were incubated with these primary antibodies overnight at 4°C on a rotator. After rinsing with PBS-Triton X-100, the discs were then incubated with the fluorophore conjugated secondary antibodies: goat anti-guinea pig (106-545-003), goat anti-mouse (115-545-003), or goat anti-rabbit (111-545-003), all purchased from Jackson Immunological Laboratories. These secondary antibodies were diluted 1:1000 in PBS-Triton X-100-NGS-BSA, and incubated with the samples for one hour at the room temperature. All samples were then stained with 1 µM DAPI at room temperature for 10 minutes, rinsed for another two times with PBS-Triton X-100 and mounted in the Vectashield mounting media (Vector Laboratories, H-1000). Confocal images were taken with a Nikon Ti Eclipse microscope system, and images were processed using the Adobe Photoshop CS6 software.

### GST-pull down assay

Full-length CDK8 fused with a N-terminal GST tag was described previously (32). The primers *mad-5.1* (F: 5’-caccATGGACACCGACGATGTGGA-3’) and *mad-3.3* (F: 5’-ctaTTAGGATACCGAACTAATTG-3’) were used for full-length Mad (AA1-455), *mad-5.1* and *mad-3.1* (F: 5’-ctaCGGGAGCACCGGACTCTCCA-3’) were used for “Mad-N1” fragment (AA1-150) that contains MH1 domain (AA10-133), *mad-5.1* and *mad-3.2* (F: 5’-ctaATCCTCCGAGGGACTGTAGG-3’) were used for the “Mad-N2” fragment (AA1-230) that contains the MH1 domain and part of the linker region, *mad-5.2* (F: 5’-caccatgCCAGTACTCGTTCCTCGCCA-3’) and *mad-3.3* were used for the “Mad-C2” fragment (AA151-455) that contains the MH2 domain (AA255-455) and part of the linker region, and *mad-5.3* (F: 5’-caccatgGGCAACTCCAACAATCCGAA-3’) and *mad-3.3* were for the “Mad-C1” fragment (AA231-455) that contains the MH2 domain. These coding sequences were amplified from a cDNA clone for *mad* gene (LD12679) using PrimeStar Max premix (Takara, R045A). The amplified products were inserted into pENTR vectors (ThermoFisher, K240020) and recombined into the pDEST17 vector (N-terminal 6XHis tag) using the Gateway LR Clonase II Enzyme mix (ThermoFisher, 11791100) in *E. coli* strain DH5α. The constructs were transformed to *E. coli* strain Rosetta for protein expression using standard protocols.

GST or GST-CDK8 was purified with Glutathione Sepharose 4B (GE Healthcare, 17-0756-01) beads with standard purification protocol. After final wash, the buffer was replaced by pull down buffer (20mM Tris-HCl pH 7.5, 10mM MgCl_2_, 100mM NaCl, 1mM DTT, 0.1% NP-40). His-tagged Mad fragments were extracted in pull-down buffer by sonication. 500µL GST or GST-CDK8 coated beads was mixed with equal volume of Mad fragments cell lysate and incubated at 4°C for 3 hours, and these samples were then washed with 1mL pull-down buffer at 4°C, 1 minute for 5 times. The interaction was detected by Western Blot with the primary antibody, anti-His (1:3000; Sigma, H1029) and the secondary antibody, anti-mouse (1:2000; Jackson Immunological Laboratories, 115-035-174).

### Yeast two-hybrid assay

Full-length CDK8 was amplified from pBS-CDK8 cDNA clone using primers *CDK8-5.1* (F: 5’-caccATGGACTACG ATTTCAAGAT-3’) and *CDK8-3.1* (F: 5’-TCAGTTGAAGCGCTGGAAGT-3’), and inserted into pENTR vector. We then used the Gateway LR Clonase II Enzyme mix to recombined CDK8 cDNA into the pGADT7-GW (prey) vector, a gift from Yuhai Cui (Addgene plasmid # 61702) (67). The linker region of Mad was amplified with *mad-5.2* and *mad-3.2* primers from a *mad* cDNA clone (LD12679) using PrimeStar Max premix, and then inserted into pENTR vector. All pENTR Mad fragments were recombined into the pGBKT7-GW (bait) vector, a gift from Yuhai Cui (Addgene plasmid # 61703) (67), using the Gateway LR Clonase II Enzyme mix. The yeast two-hybrid assay was performed using AH109 yeast strain as described previously (67).

---

**Fig. S1.**
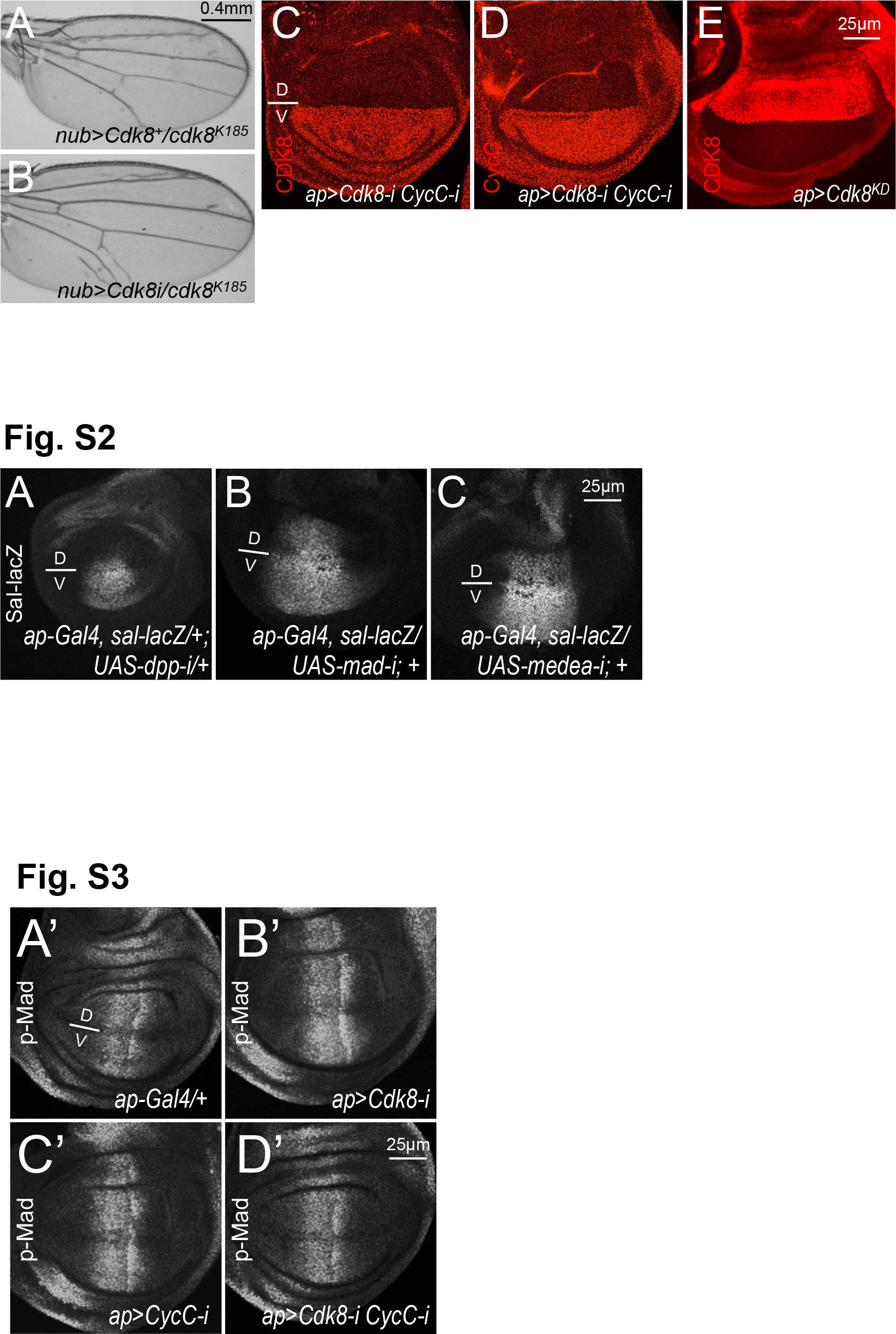
Further validation of CDK8-specific phenotypes. Adult female wings of the following genotypes: (A) *w^1118^/+; nub-Gal4>UAS-Cdk8^+^/+; cdk8^K185^/+*; and (B) *w^1118^/+; nub-Gal4/+; UAS-Cdk8-RNAi CycC-RNAi/cdk8^K185^*. Confocal images of the wing pouch area of a L3 wandering larvae wing disc of (C) *ap-Gal4/+; UAS-Cdk8-RNAi, CycC-RNAi/+* with anti-CDK8 (red) staining, (D) *ap-Gal4/+; UAS-Cdk8-RNAi, CycC-RNAi/+* with anti-CycC (red) staining and (E) *ap-Gal4/UAS-Cdk8^KD^* with anti-CDK8 (red) staining. Note that the gain in overexpression figures is lower than others, otherwise the signal will be over saturated. Scale bar in A (for A-B: 0.4mm), in E (for C-E): 25µm.

**Fig. S2.**
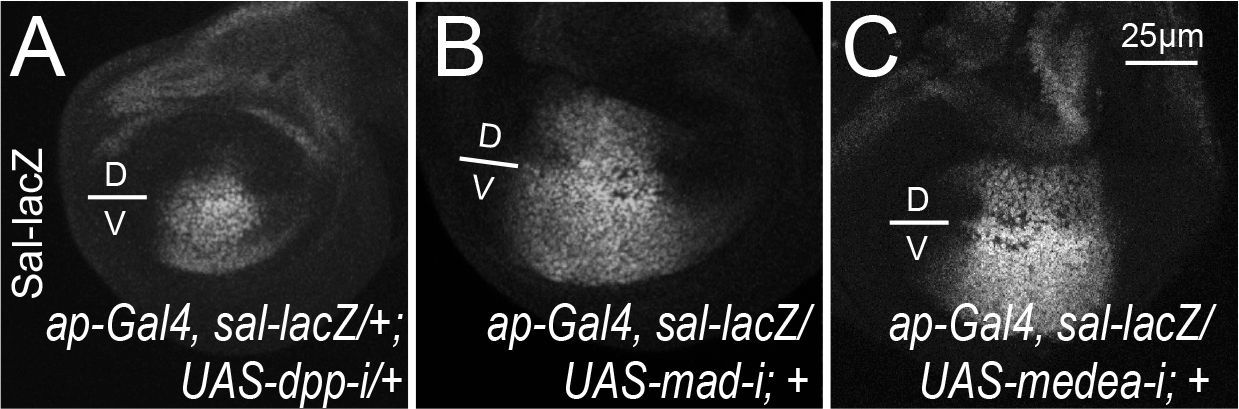
Validation of the *sal-lacZ* reporter. Confocal images of anti-β-Gal stainings of the wing ouch area of wing discs of the following genotypes: (A) *ap-Gal4, sal-lacZ/+; UAS-dpp-RNAi/+*; (B) *ap-Gal4, sal-lacZ/UAS-Mad-RNAi*; +; and (C) *ap-Gal4, sal-lacZ/UAS-Medea-RNAi; +*. Scale bar in C: 25µm.

**Fig. S3.**
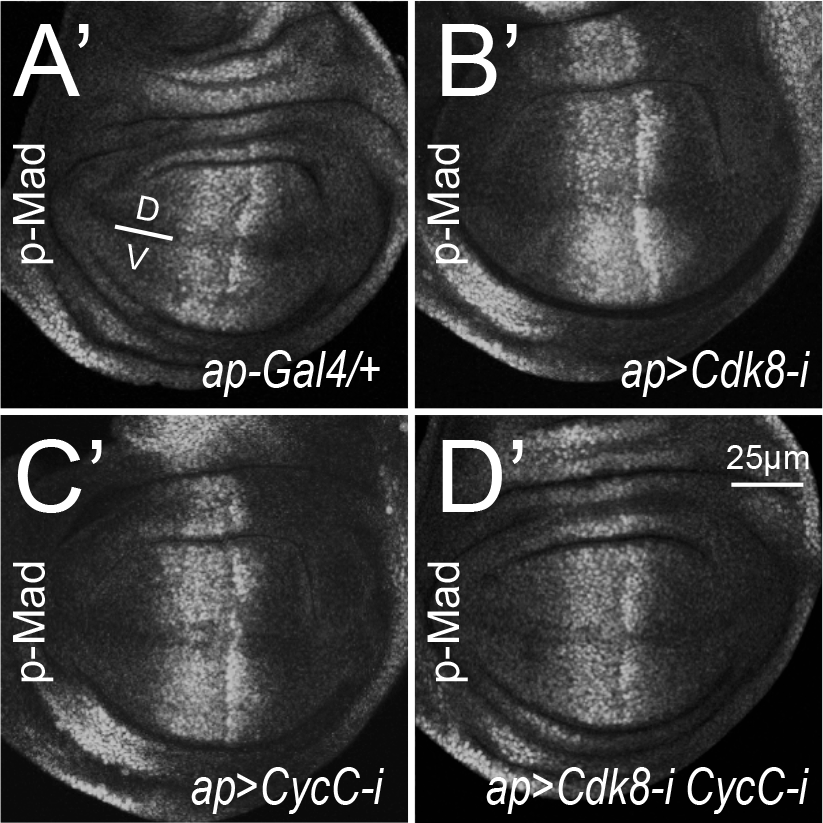
Depletion of CDK8 or CycC did not affect p-Mad level. Confocal images of anti-p-Mad staining of wing discs from the following genotypes: (A) *ap-Gal4, sal-lacZ/+* (control); (B) *ap-Gal4, sal-lacZ/+; UAS-Cdk8-i/+*; (C) *ap-Gal4, sal-lacZ/+; UAS-CycC-i*; and (D) *ap-Gal4, sal-lacZ/+; UAS-Cdk8-i CycC-i*. Scale bar in D: 25µm.

**Fig. S4.**
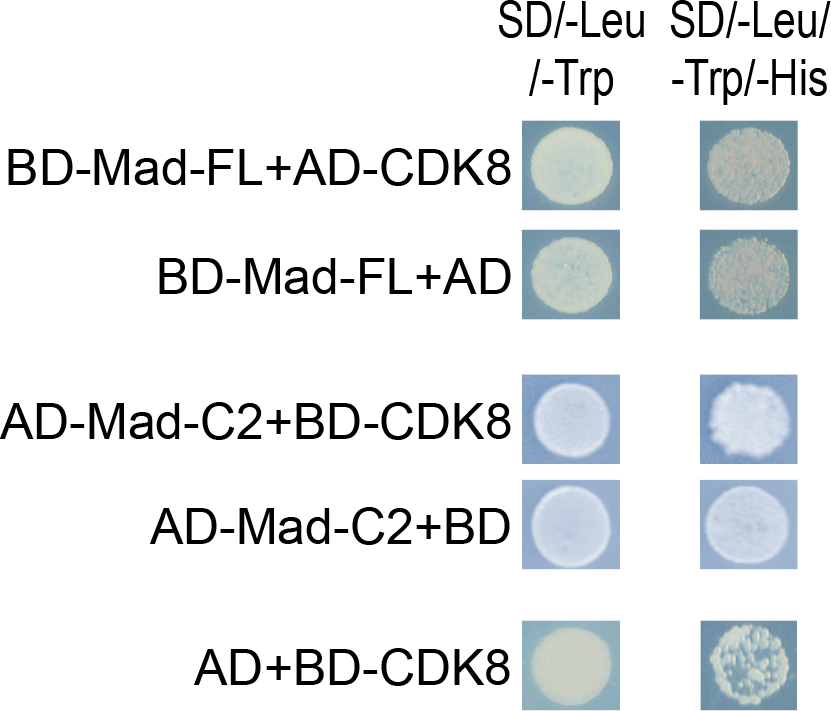
Additional results from the yeast two hybrid assay. Full-length (FL) Mad or CDK8 proteins as the bait, or Mad-C2 fragment as the prey, are able to auto-activate in this assay. Refer the figure legend in Fig. 4 and the Materials and Methods for more details.

**Fig. S5.**
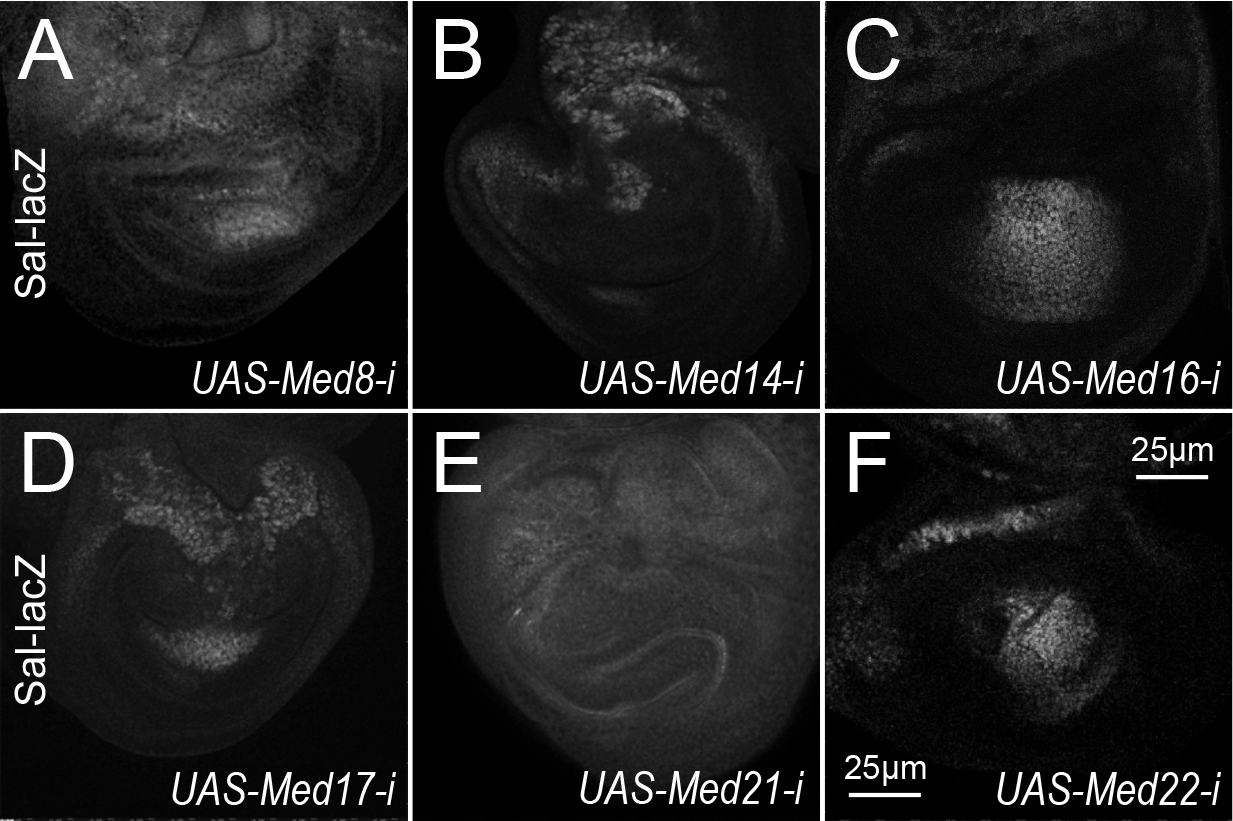
Depletion of certain Mediator subunits strongly disrupted wing disc morphology. Confocal images of anti-β-Gal staining of wing discs of the following genotypes: (A) *ap-Gal4, sal-lacZ/+; UAS-Med8 RNAi/+*; (B) *ap-Gal4, sal-lacZ/+; UAS-Med14 RNAi/+*; (C) *ap-Gal4, sal-lacZ/+; UAS-Med16 RNAi/+*; (D) *ap-Gal4, sal-lacZ/+; UAS-Med17 RNAi/+*; (E) *ap-Gal4, sal-lacZ/+; UAS-Med21 RNAi/+*; and (F) in *ap-Gal4, sal-lacZ/+; UAS-Med22 RNAi*. Scale bar in F: 25µm.

**Fig. S6.**
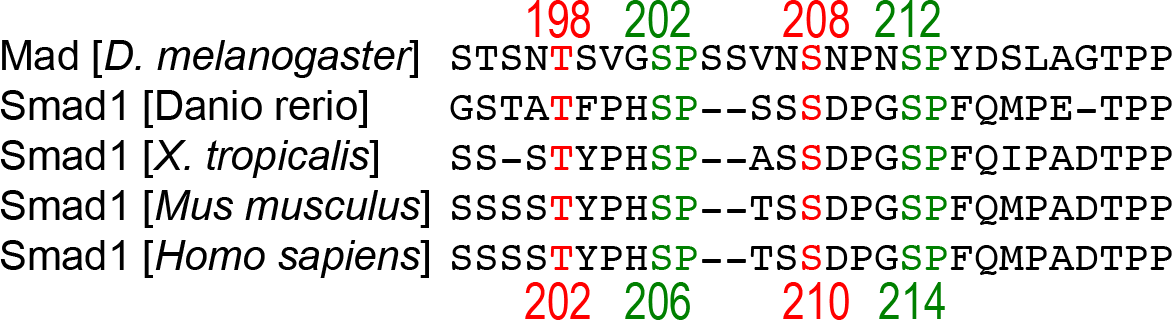
Sequence alignment of part of the Mad/Smad1 linker region showing the conservation of the potential phosphorylation sites by CDKs, MAPKs, and GSK3.

**Table S1.**
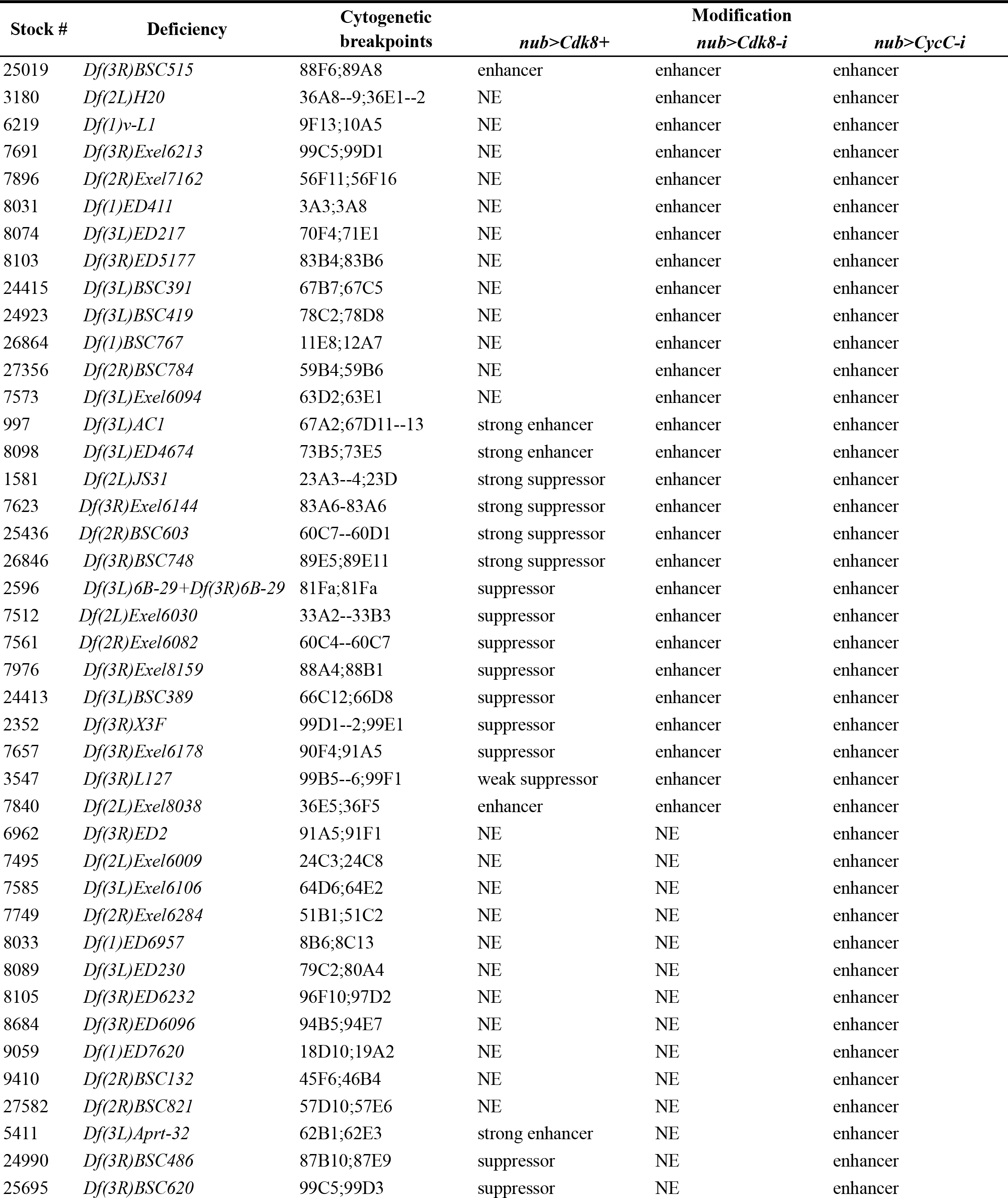
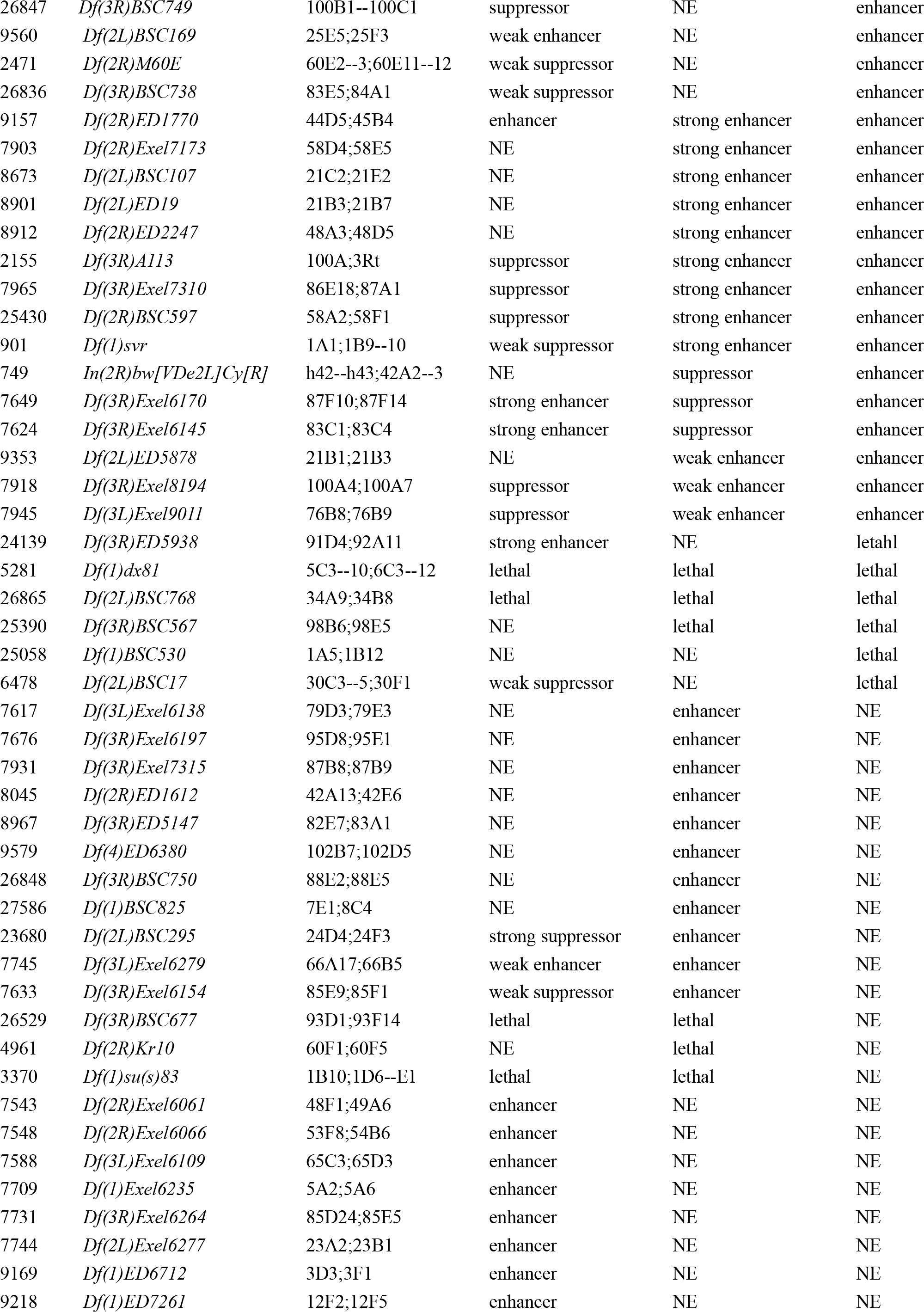
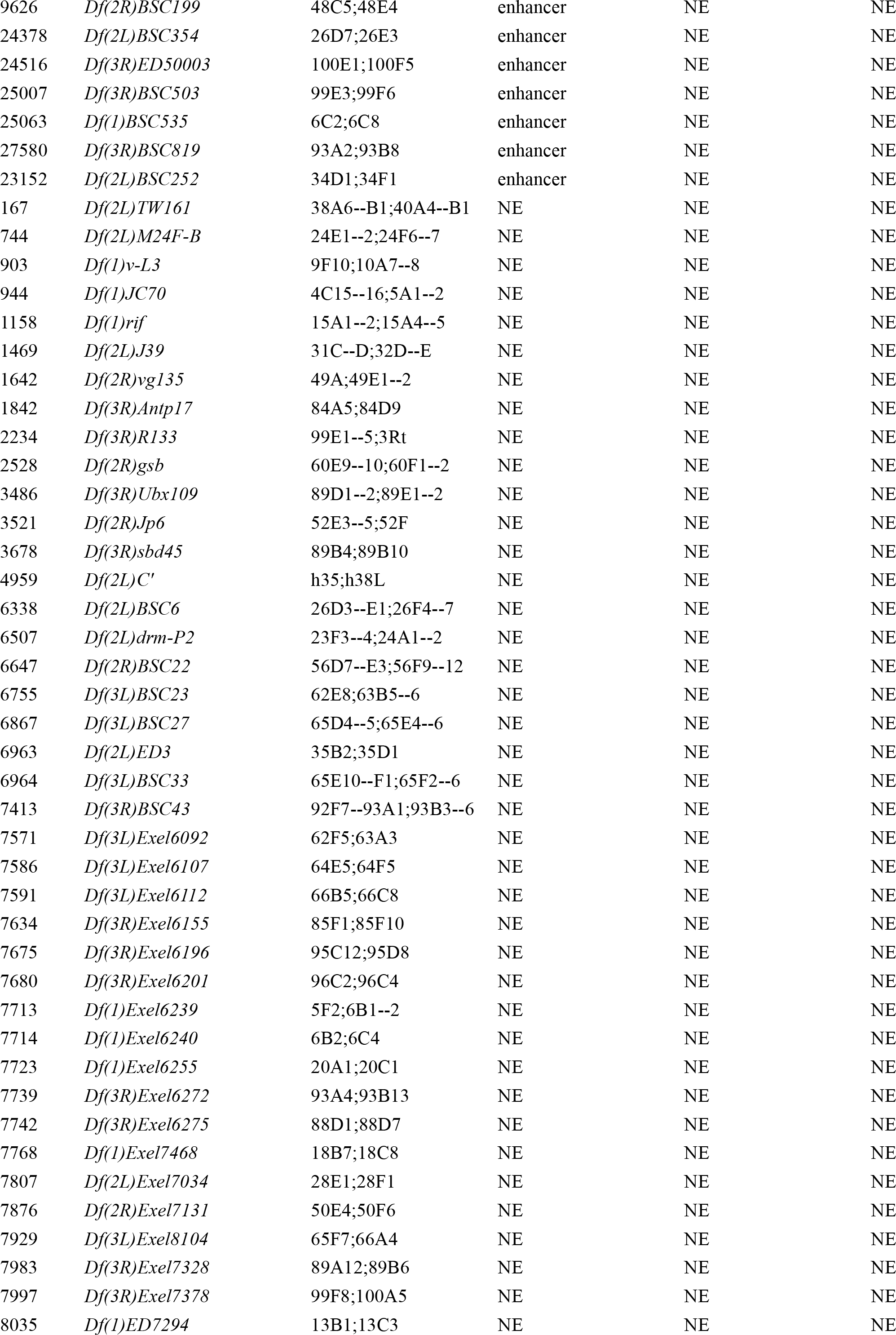
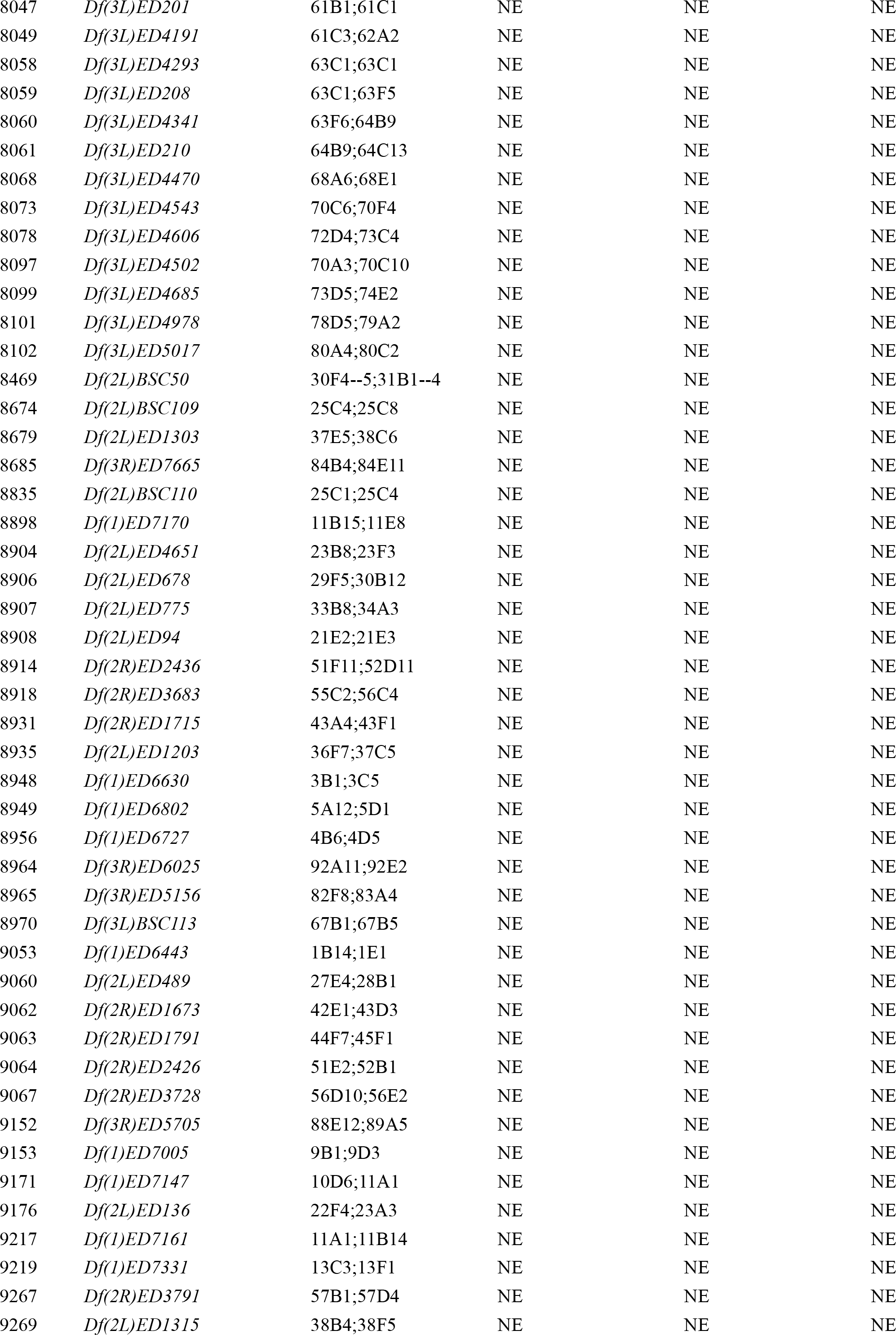
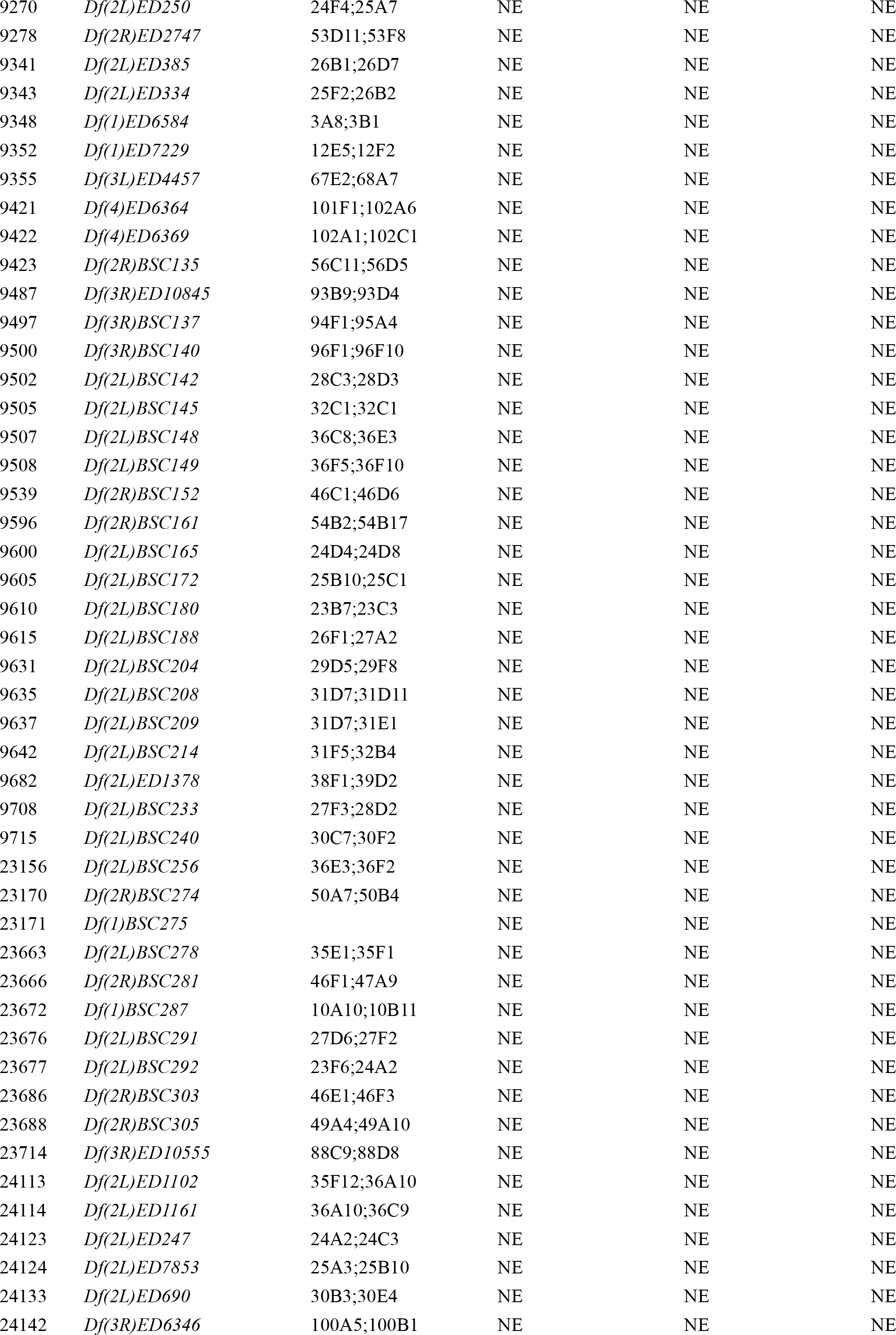
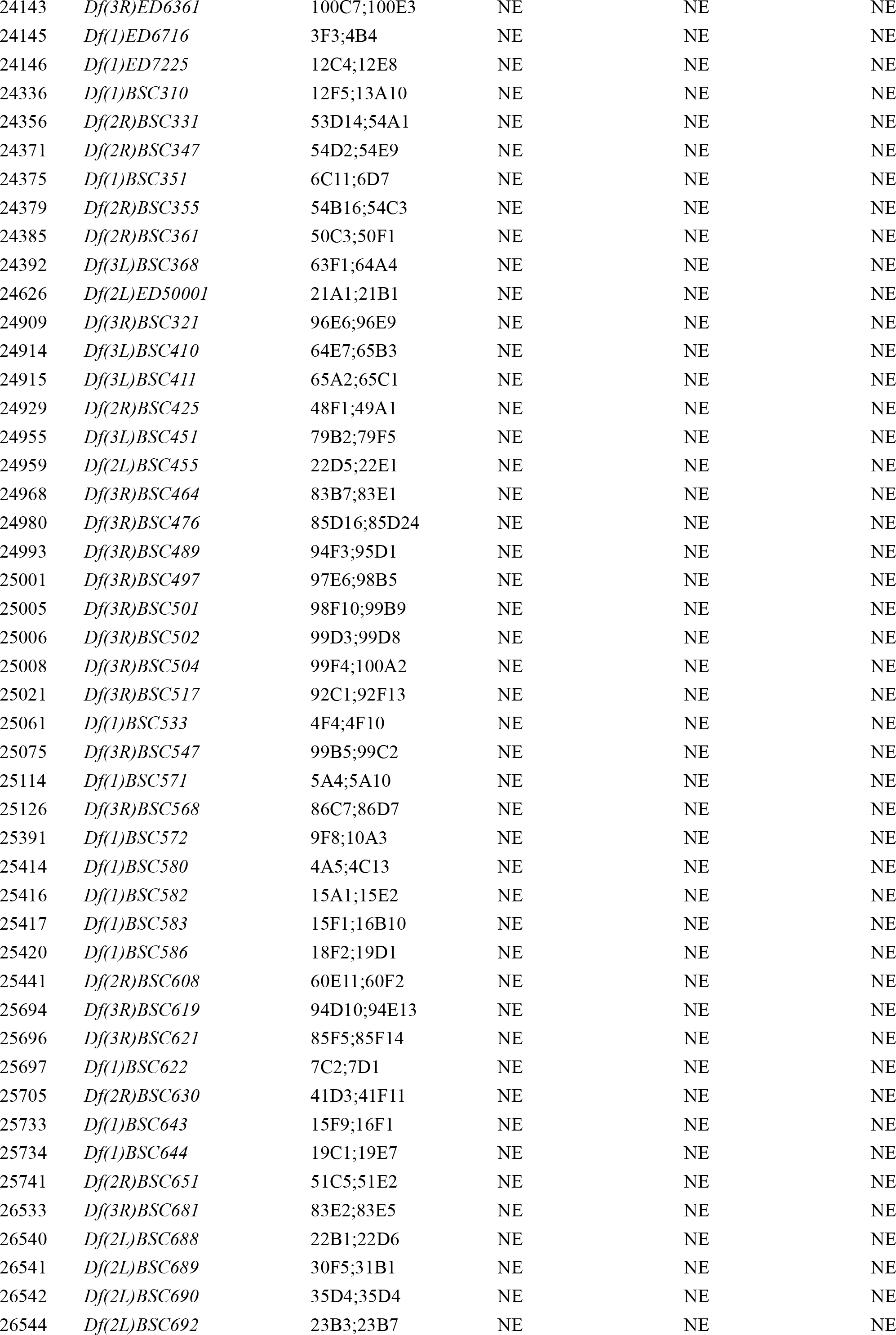
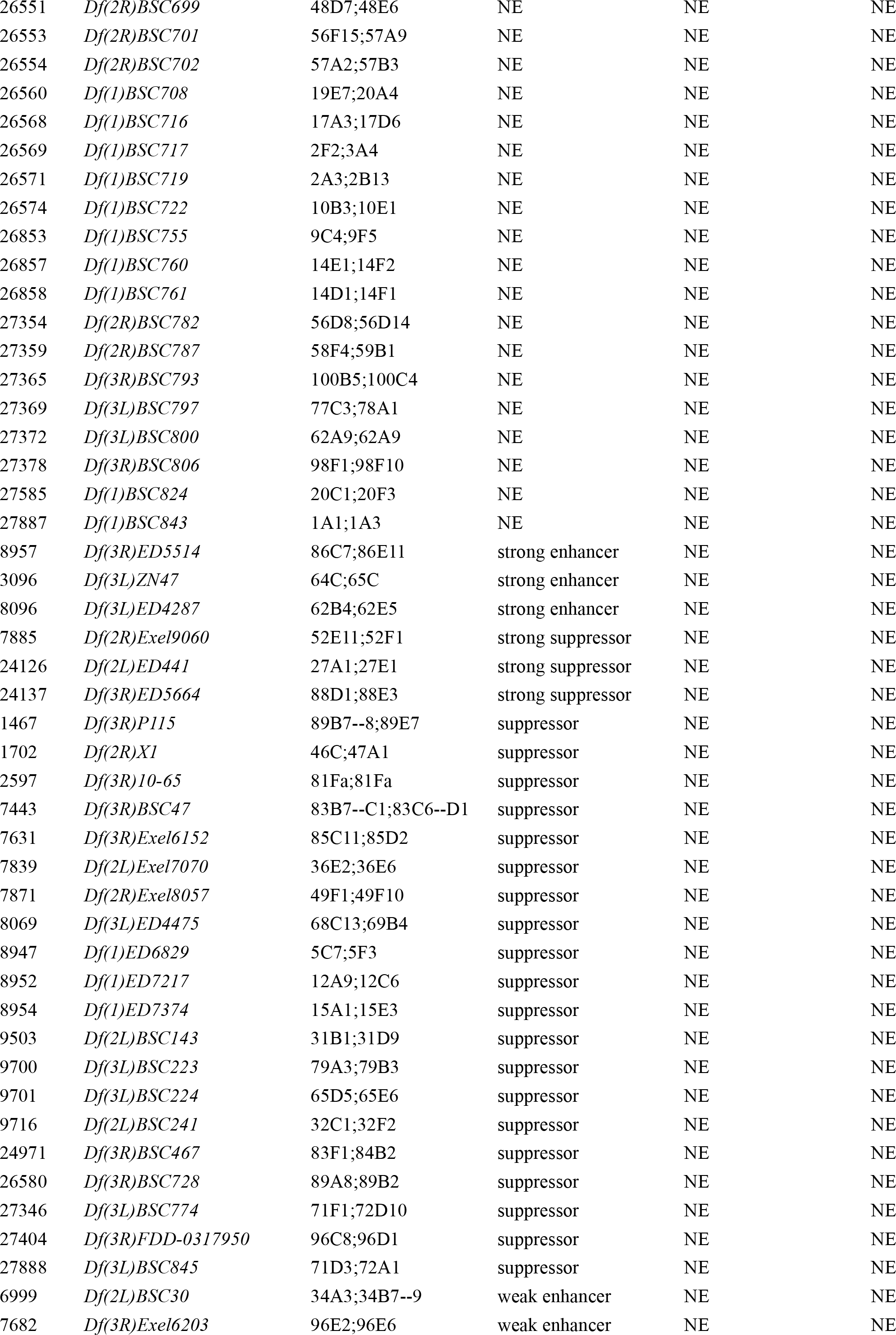
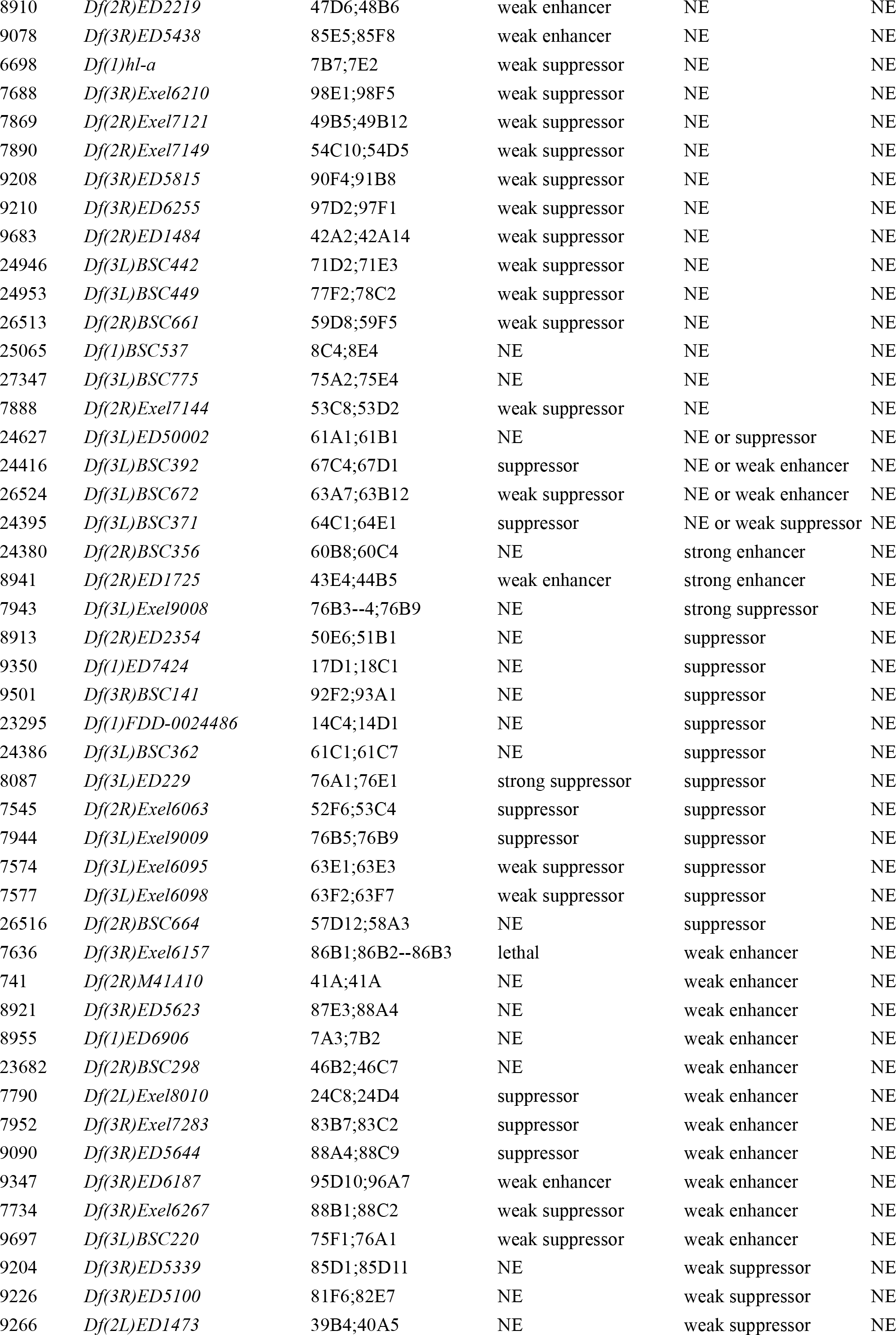
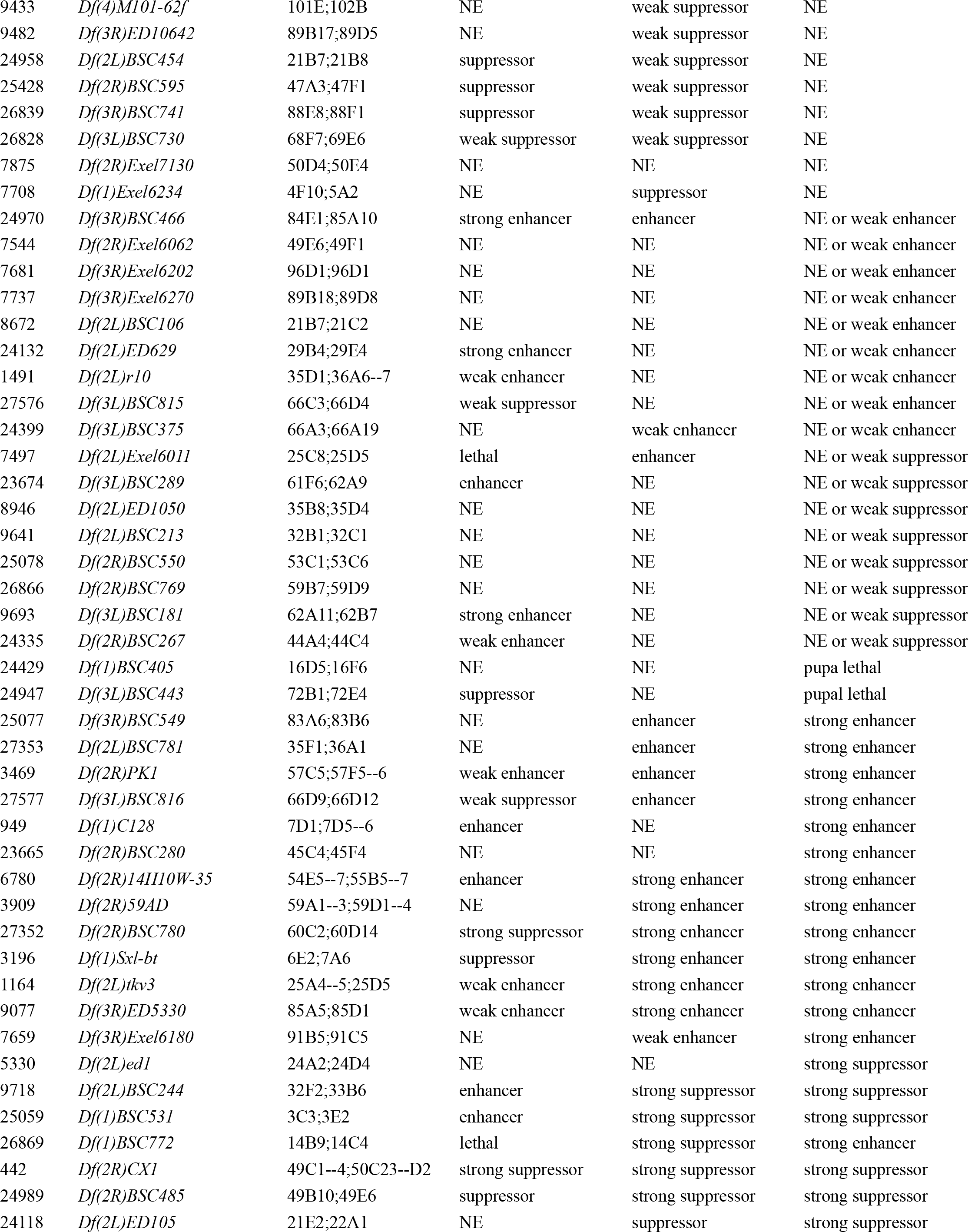
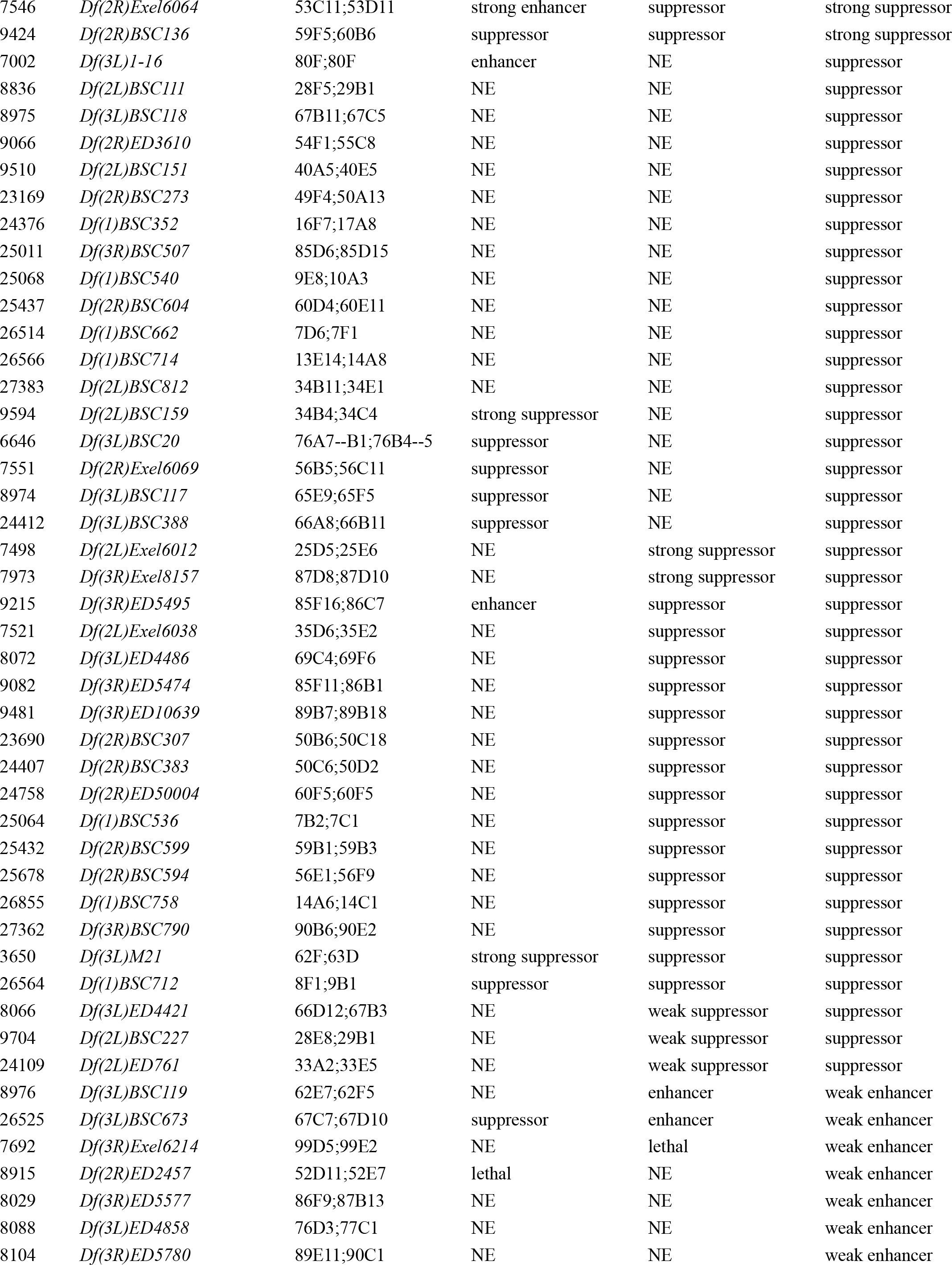
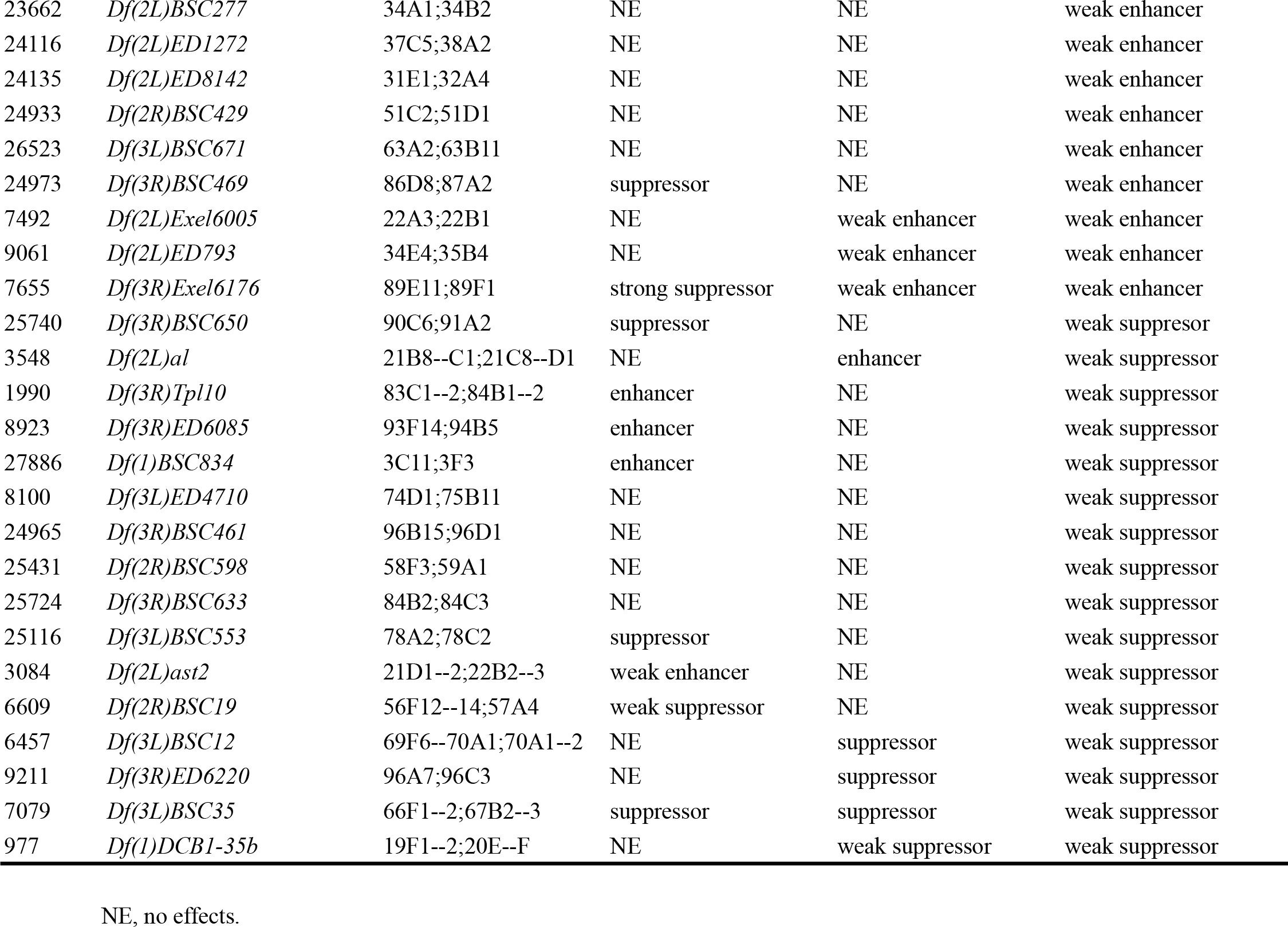
Results of 490 deficiency (*Df*) lines tested for potential dominant modification of vein phenotypes caused by altered levels of CDK8 or CycC.

**Table S2.** Specific *Df* lines that interact with CDK8-CycC.

| Stock # | Genotype | Cytogenetic breakpoints | *nub>Cdk8+* | *nub>Cdk8-i* | *nub>CycC-i* | *nub>Cdk8-i, CycC-i* |
| --- | --- | --- | --- | --- | --- | --- |
| 25059 | *Df(1)BSC531* | 3C3;3E2 | enhancer | strong suppressor | strong suppressor | enhancer |
| 901 | *Df(1)svr* | 1A1;1B9--10 | weak suppressor | strong enhancer | enhancer | lethal |
| 3196 | *Df(1)Sxl-bt* | 6E2;7A6 | suppressor | strong enhancer | strong enhancer | strong enhancer |
| 9718 | *Df(2L)BSC244* | 32F2;33B6 | enhancer | strong suppressor | strong suppressor | strong suppressor |
| 1581 | *Df(2L)JS31* | 23A3--4;23D | strong suppressor | enhancer | enhancer | enhancer |
| 25430 | *Df(2R)BSC597* | 58A2;58F1 | suppressor | strong enhancer | enhancer | weak enhancer |
| 27352 | *Df(2R)BSC780* | 60C2;60D14 | strong suppressor | strong enhancer | strong enhancer | strong enhancer |
| 7546 | *Df(2R)Exel6064* | 53C11;53D11 | strong enhancer | suppressor | strong suppressor | suppressor |
| 2596 | *Df(3L)6B-29+Df(3R)6B-29* | 81Fa;81Fa | suppressor | enhancer | enhancer | NE |
| 24413 | *Df(3L)BSC389* | 66C12;66D8 | suppressor | enhancer | enhancer | enhancer |
| 26525 | *Df(3L)BSC673* | 67C7;67D10 | suppressor | enhancer | weak enhancer | enhancer |
| 27577 | *Df(3L)BSC816* | 66D9;66D12 | weak suppressor | enhancer | strong enhancer | enhancer |
| 7945 | *Df(3L)Exel9011* | 76B8;76B9 | suppressor | weak enhancer | enhancer | enhancer |
| 2155 | *Df(3R)A113* | 100A;3Rt | suppressor | strong enhancer | enhancer | enhancer |
| 26846 | *Df(3R)BSC748* | 89E5;89E11 | strong suppressor | enhancer | enhancer | enhancer |
| 9215 | *Df(3R)ED5495* | 85F16;86C7 | enhancer | suppressor | suppressor | weak suppressor |
| 7655 | *Df(3R)Exel6176* | 89E11;89F1 | strong suppressor | weak enhancer | weak enhancer | weak enhancer |
| 7657 | *Df(3R)Exel6178* | 90F4;91A5 | suppressor | enhancer | enhancer | enhancer |
| 7965 | *Df(3R)Exel7310* | 86E18;87A1 | suppressor | strong enhancer | enhancer | enhancer |
| 7976 | *Df(3R)Exel8159* | 88A4;88B1 | suppressor | enhancer | enhancer | weak enhancer |
| 7918 | *Df(3R)Exel8194* | 100A4;100A7 | suppressor | weak enhancer | enhancer | strong enhancer |
| 2352 | *Df(3R)X3F* | 99D1--2;99E1 | suppressor | enhancer | enhancer | lethal |
| 7512 | *Df(2L)Exel6030* | 33A2--33B3 | suppressor | enhancer | enhancer | enhancer |
| 25436 | *Df(2R)BSC603* | 60C7--60D1 | strong suppressor | enhancer | enhancer | enhancer |
| 7561 | *Df(2R)Exel6082* | 60C4--60C7 | suppressor | enhancer | enhancer | enhancer |
| 7623 | *Df(3R)Exel6144* | 83A6-83B6 | strong suppressor | enhancer | enhancer | strong enhancer |

NE, no effects

**Table S3.** Mediator subunits that do not affect Mad-dependent *sal-lacZ* expression. Note that when depleted using either nub-Gal4 or ey-Gal4 line, 10 of them generated phenotypes in wing, eye, or both; while the other five are uncertain.

| **Mediator subunit** | **Effect on *sal* expression** | ***nub-Gal4* phenotype** | ***ey-Gal4* phenotype** | **Terriente-Félix et al. (2010)** |
| --- | --- | --- | --- | --- |
| Med1 | NE | Apoptosis | NE | ND |
| Med4 | NE | Vein defects | NE | ND |
| Med6 | NE | Larval lethal | small eye | ND |
| Med11 | NE | Apoptosis | ND | ND |
| Med19 | NE | Ectopic vein | ND | ND |
| Med20 | NE | Vein defects | weak defects | cell death |
| Med25 | NE | smaller wing | NE | smaller wing |
| Med27 | NE | Apoptosis | weak defects | smaller wing |
| Med28 | NE | Apoptosis | small eye | ND |
| Med30 | NE | Apoptosis | ND | cell death |
| Med9 | NE | NE | NE | ND |
| Med10 | NE | NE | NE | smaller wing |
| Med18 | NE | ND | ND | ND |
| Med26 | NE | NE | NE | ND |
| Med29 | NE | NE | ND | ND |

NE, no effects;
ND, not determined.

## Acknowledgements

We are grateful to Lauren Bridges, Christine Hermann, and Suzie Park for their technical assistance with the genetic screen, Liz Perkins for the TRiP lines, and the Bloomington *Drosophila* Stock Center (NIH P40OD018537) for the fly strains. We also thank Xiuren Zhang for advice in the yeast two-hybrid assay, and Laurel Raftery and Fajun Yang for helpful discussions. The monoclonal antibody against β-Gal (DSHB-40-1a-s) was deposited to the Developmental Studies Hybridoma Bank at the University of Iowa by Joshua Sanes. This work was supported by grants from the National Institute of Health (DK095013 to J.Y.J.) and the “Association contre la Cancer” (ARC, to H.M.B.).

